# Multiple streams of genetic diversity in Japonica rice

**DOI:** 10.1101/2020.01.23.915918

**Authors:** João D. Santos, Claire Billot, Dmytro Chebotarov, Gaëtan Droc, Mathias Lorieux, Kenneth L. McNally, Jean Christophe Glaszmann

## Abstract

In-depth studies on the genetic diversity of crops indicate that domestication is likely a drawn-out process that differs from the traditional representation of a simple rapid bottleneck. Asian cultivated rice provides a clear picture of multiple foundations of crop diversity. Among them, Japonica rice is likely the group derived from the first human manipulations of this species. We make use of the 3,000 Rice Genomes (3K RG) data set, first described in 2018, to explore the genetic diversity of traditional Japonica rice. After delineating introgressions from the Indica and *c*Aus cultivar groups, we mask these traces to analyse Japonica diversity in more depth. We find differentiation between the established “temperate”, “subtropical” and “tropical” subgroups, and identify stream-like traces of highly divergent sources from broad geographic ranges and subgroups. We characterize five such streams, most visible respectively in: 1) Indonesia, 2) continental Southeast Asia, 3) China, 4) uplands of Japan, and 5) Bhutan. These streams likely consist of ancient alien introgressions propagated through geneflow to different degrees. They currently appear as long genome segments conserved among specific germplasm groups, as well as shorter segments more broadly distributed across diverse germplasm along what could be adaptive corridors. They are all represented in the Japonica component of *c*Basmati varieties, thought to have emerged over two millennia ago. We thus provide strong evidence that Japonica, the group posited as being the most direct product of a simple domestication process in China, is an aggregate derived from multiple waves of admixture and represents a composite gene pool with ancient Asia-wide population dynamics.

## Introduction

Accurately deciphering the evolutionary history of modern staple crops is a major challenge. Compounded with previously existing genetic structure, domestication has had a major impact on the distribution of modern crop genetic diversity. *Oryza sativa,* a staple food source for over 30% of the world population, is one of the most widely distributed crops and also the first with a fully sequenced genome (Goff S. A. *et al.* 2002; IRGSP 2005). It currently benefits from an extensive sequence diversity dataset comprised of over 29 million bi-allelic markers for some 3,000 rice varieties from around the globe (3K-RG, Mansueto *et al.* 2015; Wang *et al.* 2018). This abundance of data provides an opportunity to gain greater insight into the genetic structure of modern rice.

The high phenotypic diversity observed today among *O. sativa* varieties could largely be explained by the differentiation of the two major groups, Indica and Japonica (Oka 1958). While this rift was accentuated by different domestication events and histories, it is believed to have predated the latter by more than 100,000 years, and to have taken place within the wild ancestor *O. rufipogon* (Ma and Bennetzen 2004). However, the first large-scale application of genetic markers in a study on the genetic structure of *O. sativa* (Glaszmann 1987) revealed a number of smaller groups, such as those that contain Aus-boro (*circum*-Aus, *c*Aus) or Basmati varieties (*circum*-Basmati, *c*Basmati), as well as a large number of intermediate varieties. The status of these groups as carriers of independently selected variation has received strong corroboration from their positions in the global structure analysis conducted by Wang *et al*. (2018) using the 3K-RG panel. The findings of a previous study by our group (Santos *et al*. 2019) further support these hypotheses by revealing a considerable proportion of discriminant variation among *c*Aus genomes, as well as large stretches of contiguous outlier assignments in *c*Basmati genomes. Another question brought into focus by recent results is the nature of the major groups Indica and Japonica. The prominent position of sub-populations within these groups in global clustering analyses (Wang *et al*. 2018) and the structured patterns of local classifications reported by Santos *et al*. (2019) indicate a significant degree of internal differentiation. With the *circum*-Basmati as the late product of a relatively short process of expansion and contact involving diverging evolutionary branches (Fuller *et al*. 2010), it would be plausible to expect sub-taxon differentiation to present similar patterns of incorporation into an expanding Japonica genepool.

In a recent study, we proposed a genome-wide description of a core set of landraces included in the 3K-RG (hereafter designated as CORE), based on the distribution of CORE reference groups (CRGs), representing Indica, Japonica and *c*Aus variation, at local genetic windows (Santos *et al.* 2019). This method relied on estimates of the probability density function of CRG samples in PCA feature space for the classification of local haplotypes. These were classified either as a) pure, unambiguously representing one CRG, b) as intermediate, representing a degree of uncertainty between two or more CRGs, or c) as outliers, to capture isolated and possibly cryptic material. This phenetic method allowed a detailed characterization of the CORE dataset. Despite its limited phylogenetic value and inability to correct for older genetic exchanges, this approach proved capable of roughly characterising admixed genomes, and of identifying the presence of outlier material, delineating shared material and locating recent introgressions between reference genepools (Santos *et al*. 2019). In this article, we explore research avenues opened by the genome-wide classification of local haplotypes to gain further insight into the genetic makeup of domesticated rice. This classification serves as a surface layer of information, guiding structure analyses focused on accessions and haplotypes of interest. We used the mean shift algorithm (Comaniciu and Meer 2002) to complement the supervised characterisation of local variation with an unsupervised description. The mean shift algorithm is deterministic, as pointed out by Y. Cheng (1995), and its output depends on the dimensionality of data, the number of samples and the kernels used.

We used this tool to pool information associated with the Japonica group to obtain an in-depth characterisation of the cryptic genetic variation private to this group, thus revealing its genetic makeup in greater detail than previously achieved (see Wang *et al*. 2018 for the latest structure analysis). We obtained evidence supporting the incorporation of genetic material from several differentiated sources outside of the main domesticated branches. This hypothesis is corroborated by the complementary examination of the distance this material presents to the main body of domesticated rice. Finally, we extend this analysis to the characterization of Japonica material borne by *c*Basmati genomes and consider the implications of these findings with regard to the history of Japonica rice.

## Materials and Methods

### Materials

We made use of the 3K-RG dataset generated by the 3,000 Rice Genomes Project (3K-RGP, 2014). The entire unfiltered dataset of 29 million bi-allelic markers was downloaded in November 2016. As described in Santos *et al.* 2019, one accession was removed and a missing data filter of 0.1 % was applied across markers. Analyses were run on a subset of 948 CORE landrace varieties at 137,691 overlapping windows of 150 single nucleotide polymorphisms (SNPs) along the genome (mean physical size: 5295.3 bp, sd = 9995.6). This dataset consists of 395 Indica, 320 Japonica, 65 *c*Aus, and 168 admixed accessions, among which 62 belonged to *c*Basmati (Supplemental Table S1).

### Supervised classification: kernel density estimation in PCA feature space

The kernel density estimate (KDE) of each reference group was extracted across windows. KDE was run on the first five dimensions of the projections of each CRG following principal component analysis (PCA) of the CORE dataset. The use of five dimensions complies with the recommended KDE limit (Aggarwal 2016). We find that within this limit the relations between up to eight populations are reliably captured in PCA space (Supplementary Figure S1). The function *KernelDensity* of the python package *sklearn.neighbors* (Pedregosa *et al.* 2011) was used with Gaussian kernels. CRG-specific Z scores were calculated in log-space based on the mean and standard deviation of reference accessions used for KDE only. Their lower-tail *p*-values were used to assign local haplotypes to pure, intermediate and outlier classes based on the pairwise comparison of reference *p*-values. For this we used a log *p*-value ratio threshold of 4.0 and an outlier threshold of 0.0001 (see Santos *et al*. 2019 for details).

For ideogram construction and physical summary statistics, windows were compressed by individual: in order of increasing first SNP, windows of the same classification were merged into single blocks. A new block was created with every change in class. Merged blocks range from the first SNP of the first window merged to the last SNP of the last block minus one.

### Unsupervised mean shift clustering in feature space

Following the supervised classification of genetic material at all windows considered, our objective was to describe variation specific to individual taxa in an unsupervised manner using the mean shift algorithm (MS), i.e. a density-based clustering algorithm (Comaniciu *et al*. 2002). The function *MeanShift* of the python package *sklearn.cluster* was used (Pedregosa *et al*. 2011). We used the same approach as described above for supervised classification to facilitate comparison of unsupervised clusters across datasets, i.e. 1) the KDE of each local cluster was estimated using the function *KernelDensity* function of the python package *sklearn.neighbors*; 2) the sample-level log-likelihoods of each cluster were extracted for all accessions and their lower-tail *p*-value was calculated using the mean and variance of cluster-specific observations. Cluster specific *p*-value vectors (which we refer to as MS vectors) were stored by genomic window. To recover haplotype to cluster association, each haplotype was assigned to the MS vector providing it with the highest *p*-value at locally (most affine cluster, or MAC). Haplotype-based selection of MS vectors was used for specific targeted analyses.

### Targeted analysis

We used the results of the supervised classification of local haplotypes into reference, intermediate and outlier classes, as described above. Target queries consisted of a haplotype class to germplasm taxon combination; for example, all Indica haplotypes in tropical Japonica germplasm. Here we focused on Japonica-classified haplotypes. Their MACs were used to extract an accession x cluster matrix of *p*-values and to perform PCA.

### Clustering

We first performed a classification-blind assessment as a preliminary test on the use of MS vectors for targeted local structure analysis. We performed PCA on the CORE dataset using 10,000 randomly drawn MS vectors as variables out of the total produced across every analysed window. We then performed a random but classification-specific analysis: 10,000 CRG haplotypes locally classified as Japonica were selected across all windows of consecutive SNPs where supervised analysis was conducted. The MAC of each haplotype was identified, and the corresponding MS vectors extracted (in case of coincident MACs, a single copy was extracted).

PCA was performed on the accession x MS vector matrix. Clustering could not be informed visually because of the size of the datasets generated in this step. We chose to parse this data using K-means on the transposed MS vector x accession matrix and characterize the profile of each resulting cluster individually. The *sklearn.cluster.KMeans* function (Pedregosa *et al*. 2011) was used at K=10 to generate a matrix of “affinities” (p-values) between accessions and clusters. The affinity of each accession to each of the 10 classes of local haplotype clusters was taken to be the average affinity to individual clusters within the class. The distribution of average affinities across CORE accessions by cluster was then analysed and used for discrimination of particular sub-taxa. Taking the bimodal distribution of class affinities as indicative of coherent composition, accessions were assigned to a class if their affinity was seen to lie within the second mode of this distribution. Thresholds for this assignment were visually determined.

Neighbour joining trees of local genomic diversity were constructed using the DARwin v.6 software package (Perrier & Jacquemoud-Collet 2006).

### Distance

We studied the degree of divergence that characterises the types identified through targeted analysis of MS vectors. Groups of similar MS vectors were selected for query through clustering as described in the previous section. PCA was performed on SNP data at each genomic window indexed to at least one queried vector using all CORE accessions. The first five components were kept. At each window, MAC haplotypes were recovered for each queried vector indexed to that window. Then the centroid of projections of targeted haplotypes in PCA feature space was calculated, along with the centroid of all non-target CRG haplotypes. To manage the effect of sampling, an adjusted centroid was calculated for reference CRGs as the centroid of mean shift clusters of CRG non-target accessions in feature space. We previously explored the relation between genetic correlation and Euclidian distance in PCA feature space, and it was revealed to be logarithmic and stable for a given number of markers, within an acceptable range of population number, independently of sampling (Supplementary Figure S1). Based on this latter observation, divergence was estimated as a deviation in local Euclidian distances measured in PCA space. At each window, distances were calculated to both reference and target centroids for all CORE accessions using the five captured dimensions. Deviation of individual and average distances to both centroids was measured using the mean and standard deviation of non-target CRG distances to the adjusted CRG centroid at that window. The joint behaviour of these two variables places local target haplotypes relative to both the local centre of the main body of variation and to the internal structures within it.

Two datasets were generated per query, storing vectors of target-centroid and reference-centroid distances. To establish a relation between distances at each window and the fixation index *Fst*, this latter value was predicted from raw Euclidean distance values. A linear model was used between the logarithm of *Fst* and PCA Euclidean distances adjusted to the number of SNPs used at each window. The model for each haplotype length was fitted with data generated using the simulation procedure described in Santos *et al*. (2019).

## Results

### Supervised classification of genomic windows

On average, 54.6 % of the genomes of Japonica accessions were classified in a single pure class (Table 1). Typical Japonica genotypes carried local haplotypes that could be classified as *c*Aus or Indica that on average accounted for 0.5 and 1.2% of the genome coverage, respectively. These were unevenly distributed among the Japonica subgroups identified by Wang *et al*. (2018). Tropical Japonica accessions had the largest Indica classification coverage, with 6.1 million (M) base pairs (bp) assigned to this class (q1 = 4.9 M, q3 = 9.1 M). They were followed by the subtropical (median = 3.9 M) and temperate Japonica accessions (median = 1.6 M). A similar distribution was observed for *c*Aus classifications, where the tropical Japonica group was ranked ahead of the admixed group, with an estimated median of 2.4 M bp classified in that group (q1 = 1.95 M, q3 = 2.94 M). The GJ-admx in turn had medians of 4.6 M bp assigned to Indica (q1 = 1.2 M, q3 = 7 M) and 2 M bp to *c*Aus (q = 1.4, q3 = 2.8). *Circum*-Basmati accessions presented considerably more variation in their assignment to pure classes than Japonica material, with a difference of more than 10% between minimum and maximum values across classes.

**Table 1.**
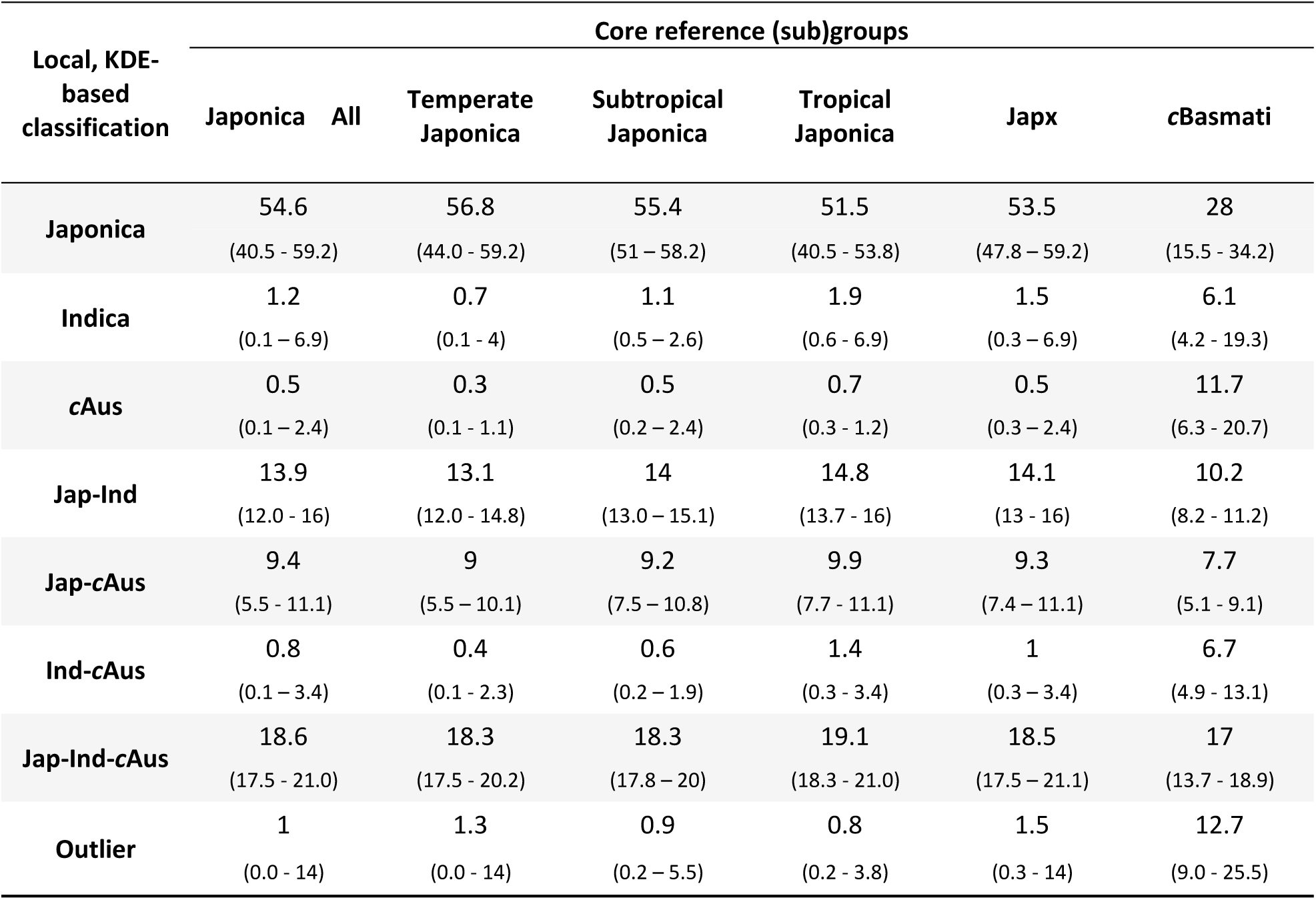
Mean percentage of genomes assigned by class using local, KDE-based classification across CORE reference groups. To estimate physical region assignment by class, local block lengths were estimated as described in Methods for supervised classification. Local block lengths were summed by class for every accession. Minimum and maximum values per group are shown in parentheses.

The remaining classes pertained to two- and three-way intermediates and to outliers. On average, local undetermined Indica-*c*Aus haplotypes represented 0.8 % of Japonica genomes, while undetermined Indica-Japonica and *c*Aus-Japonica haplotypes represented 13.9 and 9.4 %, respectively. For the other intermediate classes, which all included a Japonica component, variation was reduced among Japonica accessions and between Japonica subgroups. Among *c*Basmati material, the proportion of intermediate classes was again more variable than among Japonica material.

Outlier classifications were evenly distributed across Japonica accessions (mean = 1.0 %). Ten accessions exceeded the upper bound of this distribution range (alpha = 0.05, u.b. = 4.15 % assuming normally distributed values). Eight were temperate accessions from China, while the other two included one temperate accession from Japan (CX212, NONGKEN_58) and one subtropical accession from Thailand (IRIS_313-11848, TON TIA_GUAY NU). The outlier assignment covered a much higher proportion of *c*Basmati genomes, i.e. 12.7 % on average.

Figure 1 displays supervised classifications across tropical Japonica along chromosome 1, taken as an example. The patterns of classification reveal a diversity of situations. Long introgressions are observed as conserved discontinuities among Japanese and South Korean samples with one Chinese sample at the bottom of the list. Here, extended stretches of pure or intermediate Indica classifications suggest introgressions, reaching as long as 27 Mb in more than one case. However, the presence throughout this region of smaller structural gaps, particularly involving Japonica sources, hampered accurate determination of their limits. Besides this feature, assignment alternations can appear as isolated events, restricted to a few accessions, as well as shared events common to a few or many accessions. In the latter case, shared haplotypes were classified as ambiguous between Japonica and either Indica (5-10 Mb), *c*Aus (5-10 Mb) or both (20-21 Mb), and they appeared as vertical non-blue columns. Such broadly conserved discontinuities are likely to represent extensive genetic diffusion events across rice groups. Less conserved patterns denote more limited exchanges and result in apparent insertions in Japonica genomes of haplotypes assigned outside Japonica. This could be seen as a distribution of Indica material in several regions, notably at 13-14 Mb, 23-24 Mb and 28-30 Mb. Alternatively, variations in discontinuity patterns between shared and pure Japonica classifications, as noted within 1-3 Mb, indicates the co-existence of differentiated material within this subgroup. Isolated introgressions appear clearly delineated, notably contiguous *c*Aus assignments within 4-8 Mb and 20-23 Mb regions. Outlier classifications appear to be only slightly structured, with no indication of local alien introgressions of significant size.

**Fig. 1.**
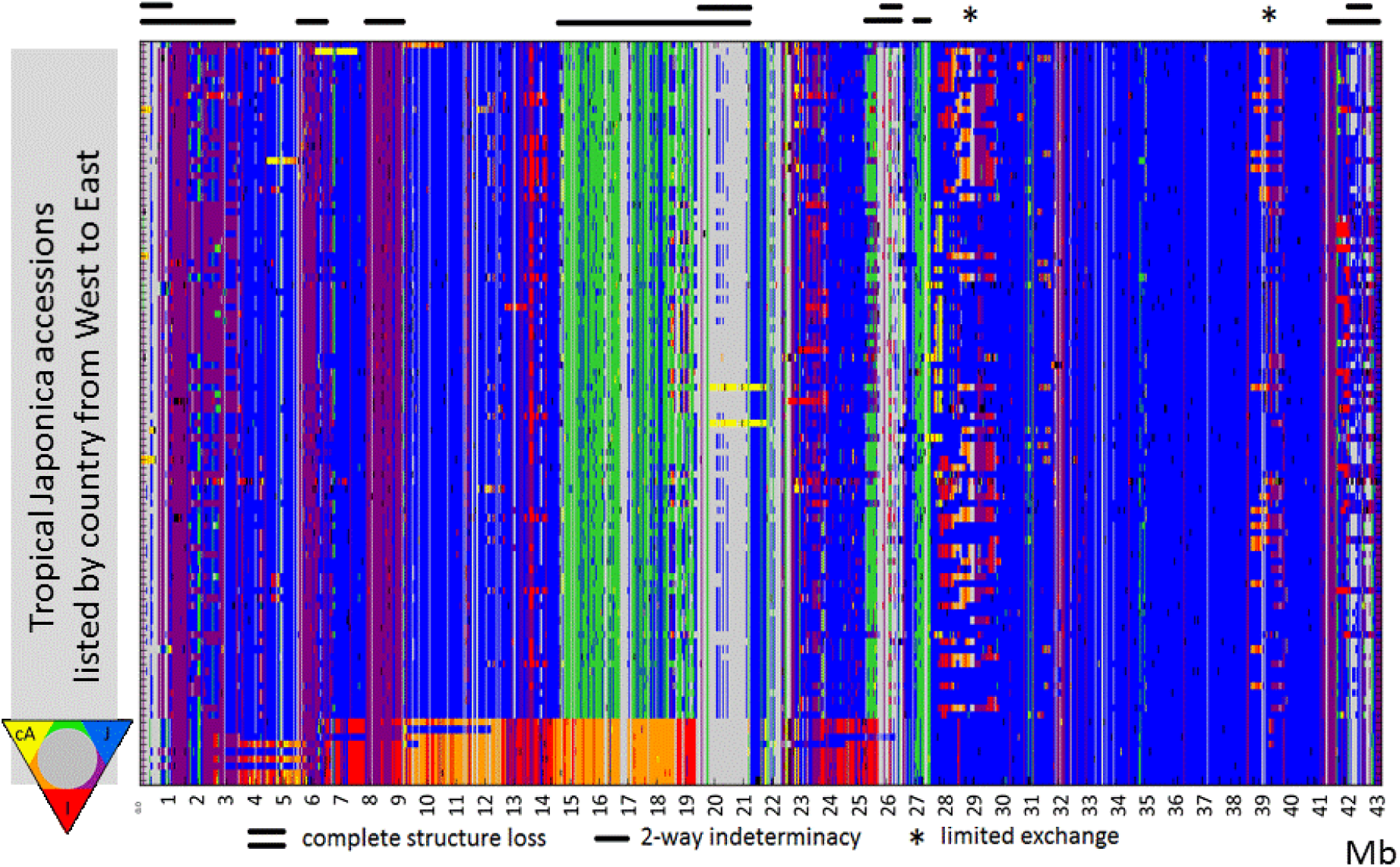
Ideogram of a supervised classification of local haplotypes across tropical Japonica chromosome 1. Individual accessions are ordered vertically on a geographical basis, and genome positions horizontally. Local haplotypes are classified based on pairwise comparison of reference population *p*-values. *P*-values were extracted as normalized KDE likelihood estimates. Genome assignment colour codes are given in the bottom left triangle; black = outlier.

### Global and Japonica variation – MAC-mediated exploration

The local haplotype cluster affinity matrix (MS vectors, see Methods) resulting from the window-based analyses recovered the classification of Wang *et al*. (2018) and highlighted deeper substructural elements in each group. The classification-blind PCA run based on 10,000 randomly selected local clusters captured the three major rice groups, i.e. Indica, Japonica and *cAus*, within the first two PCs (Fig. 2A). The third principal component revealed the Japonica substructure (Fig. 2B). In the Japonica-targeted analysis based on 10,000 randomly selected clusters, the first two PCs were in line with the most recent temperate, subtropical and tropical subgroup classification for this group.

**Fig. 2.**
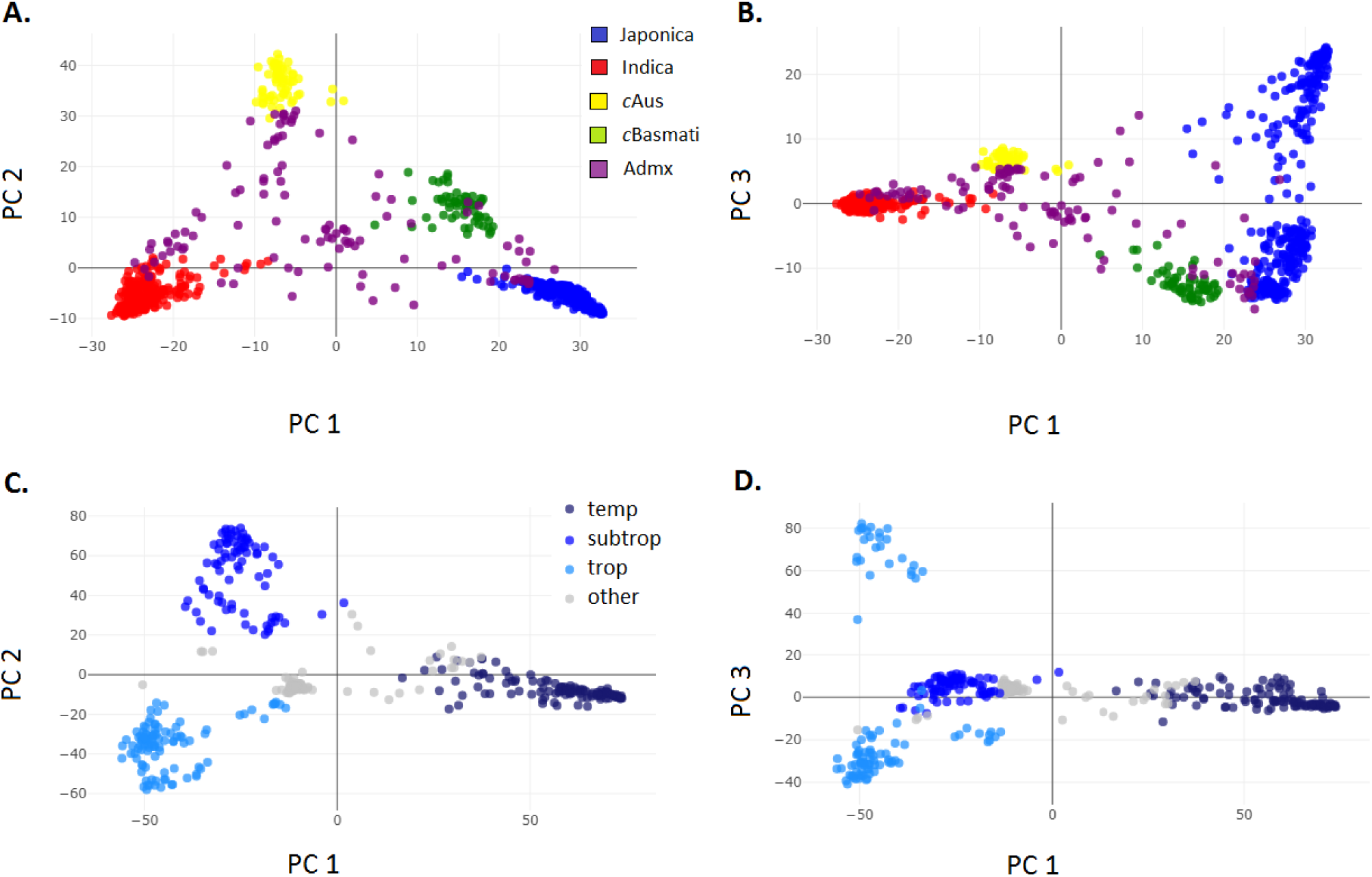
Non-target and reference specific associations. PCA of CRG accessions using mean shift vector affinities. **A**-**B**) first three PCs of the PCA on random MS vectors. Global sNMF groups are displayed. PC var. explained: I: 11.8 %, II: 6.2 %, III: 2.9 %; **C**-**D**) first three PCs of the PCA on Japonica targeted MS vectors. Japonica subgroups of Wang *et al*. (2018) displayed. PC var. explained: I: 11.8 %, II: 7.1 %, III: 3.7 %.

The first PC separated temperate Japonica, whilst the second highlighted the subtropical *vs*. tropical Japonica variation (Fig. 2C). Subgroup-specific correlations were captured in the dimensions beyond the first two PCs, with the third principal component isolating (on its positive side) a subgroup of 26 Indonesian tropical Japonica accessions (Fig. 2D).

### Japonica sub-structure

The MACs of Japonica classified haplotypes across CORE genomes were queried, returning 290,756 MS vectors. We studied projections of CORE accessions following PCA of the Japonica variation matrix (Japonica accessions *x* Japonica selected MS vectors).

Based on our analysis, local haplotypes were assigned to ten classes and the affinity to each of these classes was estimated for each of the accessions. At K = 10, the number of vectors per group varied widely, with six groups representing less than 8 % of the total number of vectors each, while one group represented 41 % (see Figure S2). We used the distribution of affinities per class to assign accessions to subgroups containing accessions most affine to the corresponding class, and specified a threshold for each class that best reflected the shape of the likelihood distribution, which was usually clearly bimodal (Figures S2 to S4, Table S1). Ultimately, we retained eight subgroups that consisted of at least five accessions that were markedly more affine to their cluster than to any other. A ninth subgroup (corresponding to class 2) consisted of only two accessions from China and Korea. The affinity distribution to class 1, i.e. the group with the largest MS vector number, did not enable us to identify a specific subgroup, indicating that it was a haplotype class common to all Japonica accessions.

We used the above-mentioned PCA planes to visualize the distributions of these subgroups and thus help clarify their nature (Fig. 3).

**Fig. 3.**
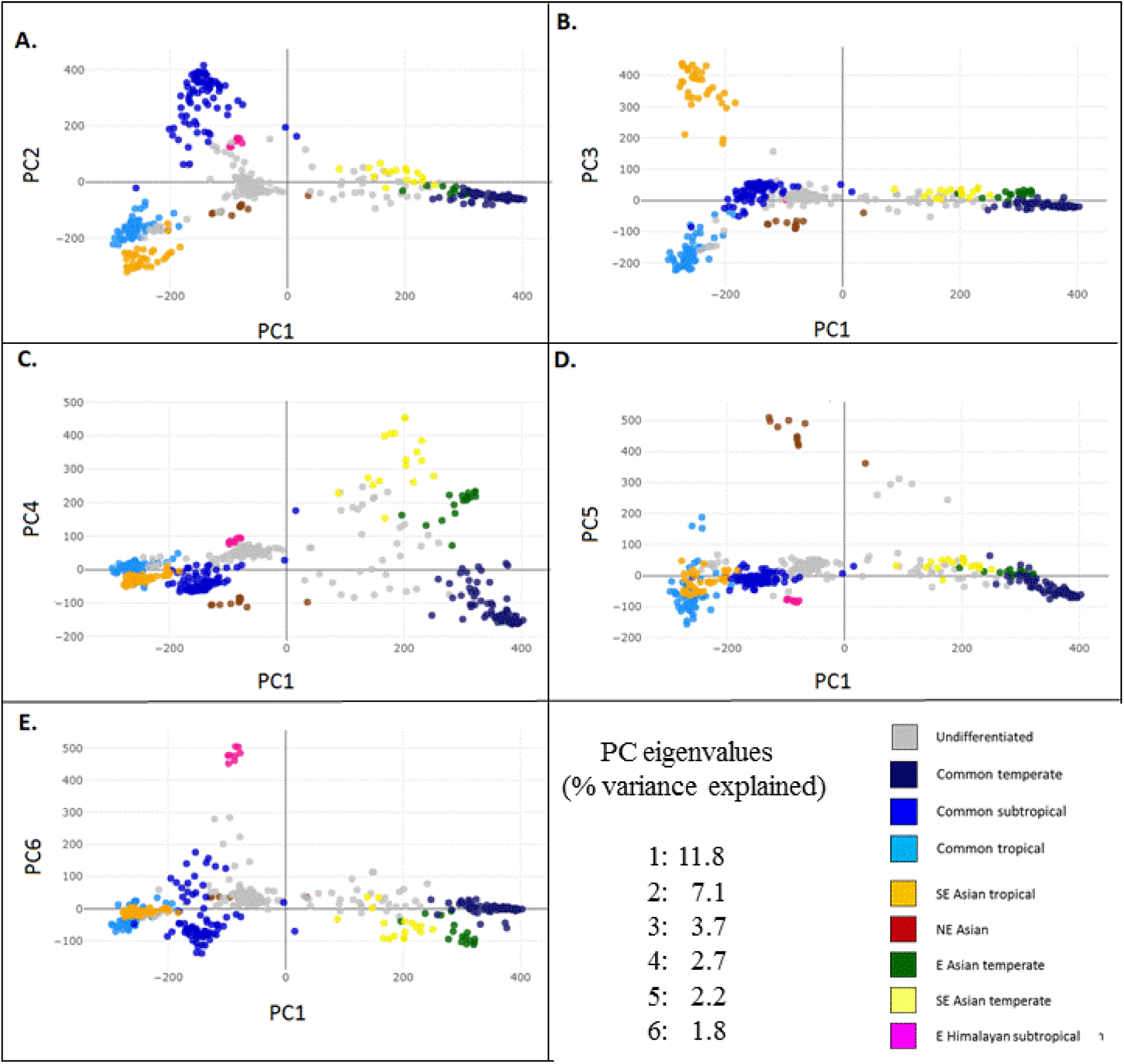
Deep structure of the Japonica complex. Principal component analysis using mean shift vectors of Japonica MACs across major Japonica subgroups. **A**) PCs 1 and 2. **B**) PCs 1 and 3. **C**) PCs 1 and 4. **D**) PCs 1 and 5. **E**) PCs 1 and 6. Subgroup assignments were determined using a lower threshold on subgroup-specific average affinities. Lower thresholds were manually established by visual inspection of subgroup-specific distributions (Supplementary Figs. 6, 7, 8).

We combined likelihood profiles and geographic distributions and summarized the new subgroup landscape as follows:

- a “common temperate type” (cJtm) (corresponding to class 8 in Table S1), visible along axis 1, pooling accessions from various essentially East Asian origins, but also from western Asia, classified in the temperate subgroup of Wang *et al.* (2018).
- a “common tropical type” (cJtr) (class 6 in Table S1), visible along axis 3, pooling accessions from various in Malaysian, Indonesian and Philippine origins, often grown in upland environments, classified in the tropical subgroup of Wang *et al.* (2018).
- a “common subtropical type” (cJsb) (class 7 in Table S1), visible along axis 2, pooling accessions from various continental Southeast Asian and Northeast Indian origins, often grown in upland environments, classified in the subtropical subgroup of Wang *et al*. (2018).
- a “Southeast Asian tropical type” (SEA-Jtr) (class 9 in Table S1), symmetrical to subgroup 6, visible along axis 3, essentially pooling accessions from Indonesia, classified in the tropical subgroup of Wang *et al*. (2018).
- a “Southeast Asian temperate type” (SEA-Jtm) (class 4 in Table S1), visible along axis 4, pooling varieties from Vietnam, Malaysia, the Philippines and China and classified, despite their distributions, in the temperate subgroup or as Japonica-admixed of Wang *et al.* (2018).
- an “East Asian temperate type” (EA-Jtm) (class 3 in Table S1), visible along axes 4, 9 and 10, essentially pooling varieties from China, classified in the temperate subgroup of Wang *et al.* (2018).
- a “Northeast Asian type” (NEA-J) (class 10 in Table S1), visible along axis 5, pooling mostly upland varieties from Japan and Korea, but yet classified in the tropical subgroup of Wang *et al.* (2018).
- an “East Himalayan subtropical type” (EH-Jsb) (class 5 in Table S1), visible along axis 6, pooling varieties mostly from Bhutan, classified in the subtropical subgroup of Wang *et al.* (2018).

Table 2 summarizes accession genome compositions averaged by class as defined in Table S1 (MS column). Based on the assumption that window assignments to Class1 indicated a lack of specificity, other assignments depicted the true specificities. The highest values in the columns are highlighted. Four subgroups had a major share of their specific genome assigned to themselves: cJtm, cJsb, cJtr and SEA-Jtr. All the others had the largest part of their genome assigned to cJtm. On average, the most frequent assignments were to cJtm, followed by cJsb, then cJtr and NEA-J.

**Table 2.** Average genome proportions (per thousand) assigned to classes of Japonica sub-specificity within each subgroup. To estimate the physical region assignment by class, local block lengths were estimated as described in Methods for supervised classification. Local block lengths were summed by class per accession. The highest average values in the columns are highlighted

| Haplotype class | Accession group |  |  |  |  |  |  |  |
| --- | --- | --- | --- | --- | --- | --- | --- | --- |
|  | cJtm | cJsb | cJtr | SEA-Jtr | SEA-Jtm | EA-Jtm | NEA-J | EH-Jsb |
| Class 1 | 515 | 537 | 606 | 550 | 496 | 490 | 541 | 553 |
| Class 2 | 21 | 19 | 15 | 16 | 22 | 24 | 18 | 19 |
| **cJtm** | 445 | 145 | 69 | 93 | 287 | 347 | 142 | 158 |
| **cJsb** | 35 | 226 | 61 | 54 | 56 | 45 | 79 | 90 |
| **cJtr** | 12 | 35 | 115 | 61 | 14 | 12 | 23 | 30 |
| **SEA-Jtr** | 5 | 7 | 20 | 131 | 7 | 7 | 13 | 7 |
| **SEA-Jtm** | 5 | 6 | 4 | 0 | 53 | 12 | 3 | 5 |
| **EA-Jtm** | 17 | 7 | 5 | 6 | 36 | 87 | 4 | 11 |
| **NEA-J** | 14 | 20 | 37 | 31 | 12 | 13 | 116 | 13 |
| **EH-Jsb** | 7 | 24 | 15 | 10 | 16 | 9 | 8 | 103 |

### Fine Japonica genomic differentiation

The affinity of haplotypes to target MACs was plotted against accession genomes in the form of ideograms. These patterns can display patches of contrasting assignments characterized by their breadth (frequency within the taxon under consideration), continuity (assignment homogeneity along the segment) and length (distance between the start and end of the continuous segment).

The genome-wide patterns of MS subgroup-classified local haplotypes shown in Table 2 are outlined in Figure 4 and Supplementary Figures S5 through S7. These figures show the distribution of the classified haplotypes identified by specific colours, zones of lack of assignment in grey, and zones assigned previously as intermediate, outlier, pure Indica or *c*Aus in white. Many coloured patches are short and dispersed. However, some have a high level of continuity over a significant length and breadth, as can be seen on chromosomes 3 and 6 to 10 (Figure 4).

**Fig. 4.**
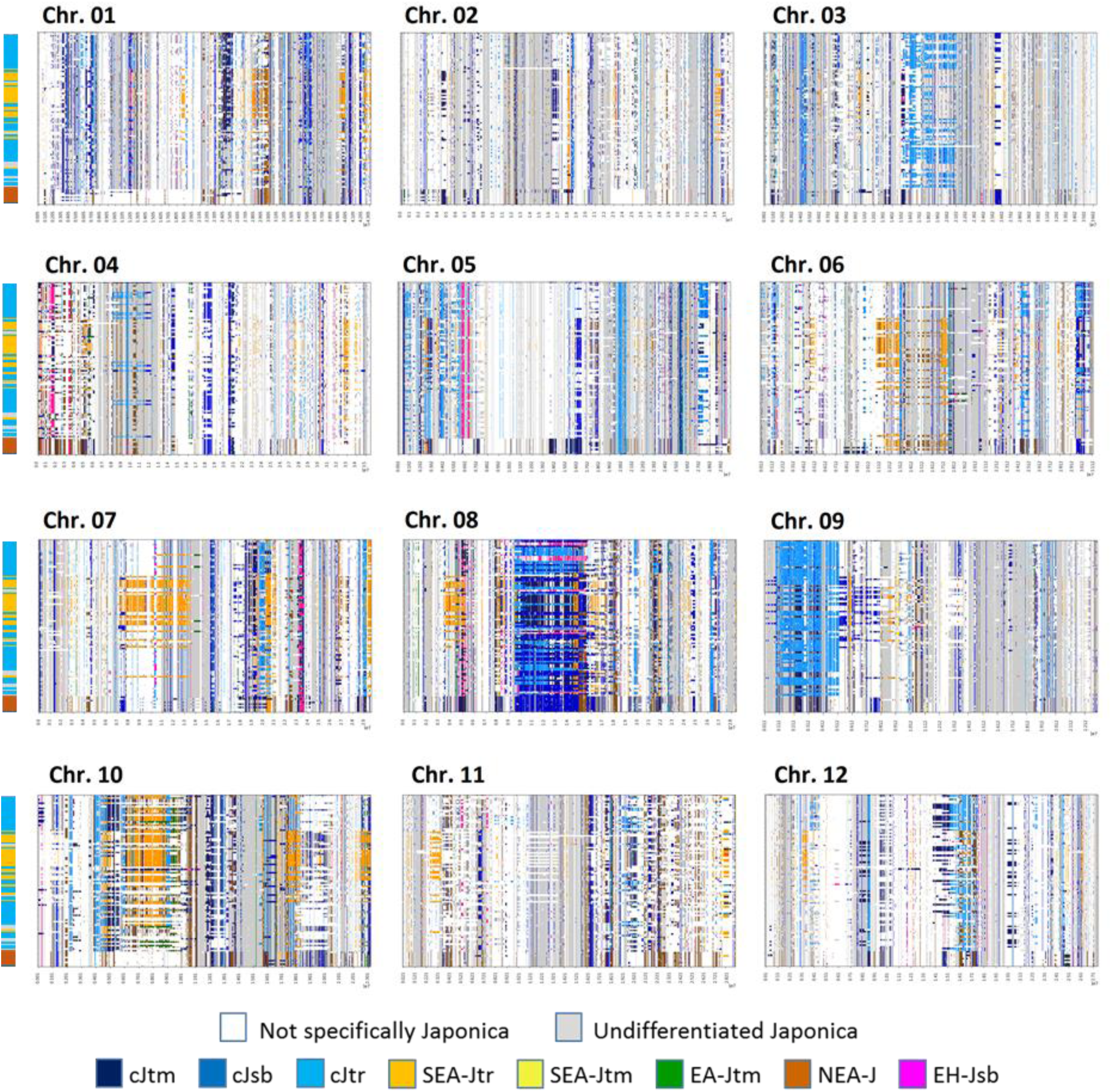
MAC assignment to subgroups indicative of the Japonica substructure along the genome of tropical Japonica accessions. Local cluster vector *p*-values were extracted for pure Japonica assignments across Japonica genomes. Vectors were grouped following PCA using k-means clustering at K = 10. The vertical box on the left shows the classification given in Table S1. Local tropical Japonica haplotypes were analysed for maximum affinity to clustered vectors and classified accordingly.

Overall, a comparison of the three major established subgroups highlighted the broader distribution of haplotypes associated with the common temperate class (dark blue), with comprehensive genome coverage in this subgroup and a frequent occurrence in common tropical and subtropical subgroups. Haplotypes associated with the common subtropical class (blue) had the second broadest distribution, as was also the case in the tropical subgroup but not the temperate subgroup. Figure 4, which describes accessions classified in the tropical subgroup, shows many patches related to common temperate and subtropical classes as well as other more specific patches initially associated with the corresponding k-means subgroups observed in Figure 3. The latter could have distributions restricted to their corresponding class (Supplementary Figures S5 through S7), as on chromosomes 1 and 8 for Southeast Asian tropical (orange) signals (Fig. 5), or beyond the originally perceived coherence, as on chromosomes 6 or 10, for signals extending further within the same subgroup (orange). We selected chromosome 8 to display patterns for all Japonica accessions in Figure 5 (we include *c*Basmati for further discussion below).

**Fig. 5.**
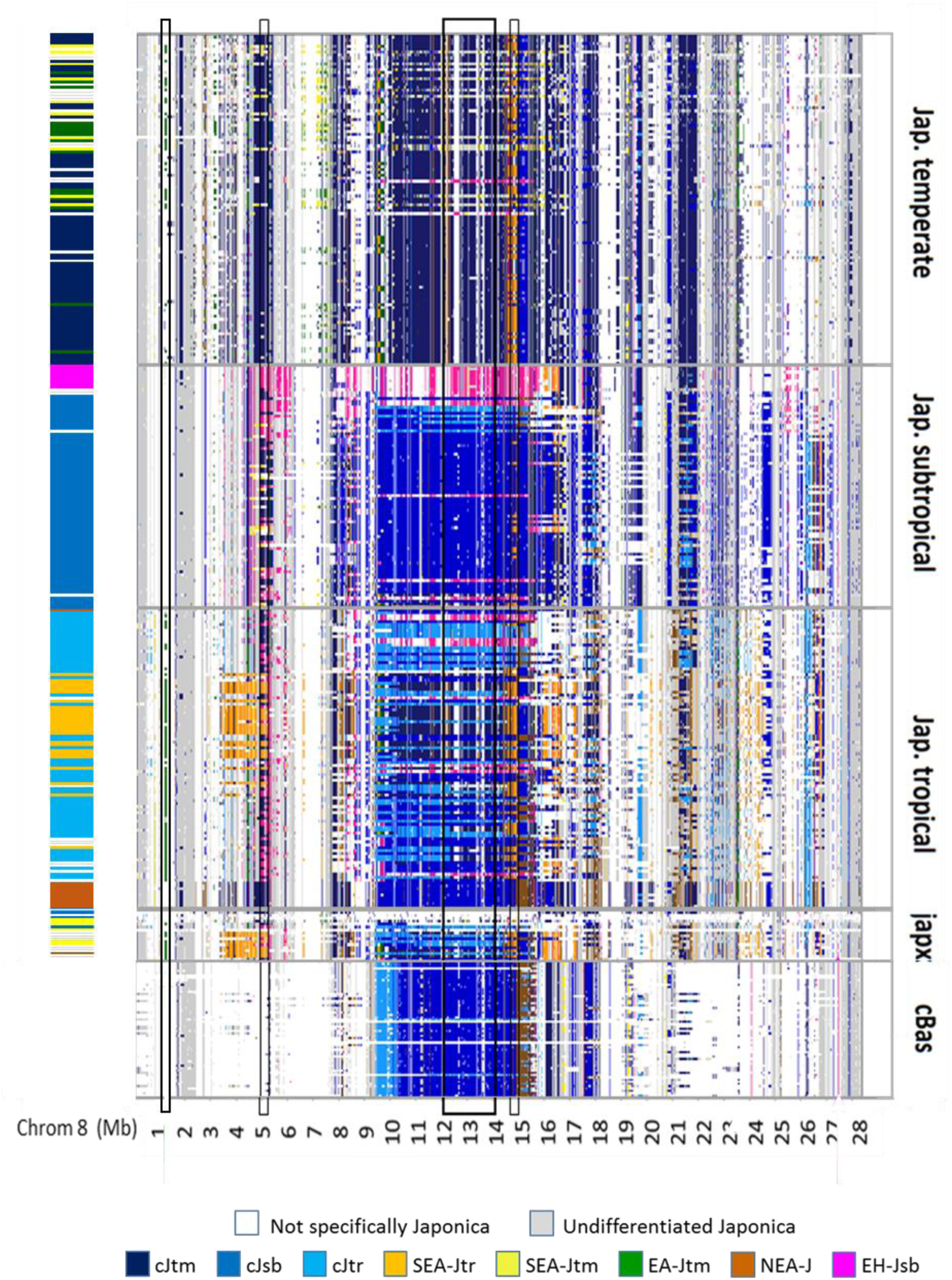
Japonica sub-taxon affinity along chromosome 8 as predicted by MS vector analysis. Japonica classification was used to extract MAC vectors across Japonica subgroups and the *c*Basmati. At each query, vectors were grouped using k-means following PCA, and the group mean *p*-value profile was analysed. Average profiles indicative of Japonica substructure were selected and the local haplotypes involved retraced and depicted accordingly. The block on the left indicates global assignment to Japonica subgroups. The bottom part features *c*Basmati accessions to illustrate their Japonica component (see below). Vertical boxes represent the segments used for more detailed analysis. Four regions were identified and used in Figure 6 for constructing NJ trees.

On chromosome 8 (Figure 5), around the central portion of the chromosome (third identified region), we found tropical haplotypes assigned to the EH-Jsb class (pink). This variation in assignment extending across the three Japonica subgroups was clearly visible when focusing on the central chromosome regions, with tropical accessions bearing haplotypes assigned to temperate and subtropical discriminant clusters and subtropical accessions bearing haplotypes assigned to temperate clusters (dark blue). The distribution of haplotype assignments to different classes across subgroups is suggestive of streams wherein blocks of historical introgression from distinct sources achieved different degrees of penetration through geneflow between populations. We relied on this representation to select a few instances to monitor the extent of differentiation along the streams.

### Local structure along introgressive streams

In order to validate the differentiation revealed with our method and to assess the respective levels of divergence, we constructed Neighbour-joining trees of local genomic regions where the MAC-mediated assignment of local haplotypes displayed specific patterns when considering all accessions of the CORE set. We chose four regions on chromosome 8 (Fig. 5) that represented different patterns. Some were continuous stretches of variation discriminant of the sub-taxa queried suggestive of long introgression tracts, while others were smaller segments attached to a specific subgroup that had a broader distribution than the subgroup itself, suggestive of positive selection. The resulting NJ trees are shown in Figure 6 in order of increasing genomic coordinates. We found that this classification method effectively revealed the differentiated branches of genetic diversity corresponding to our queries (Figures 4, S6, S7).

**Figure 6.**
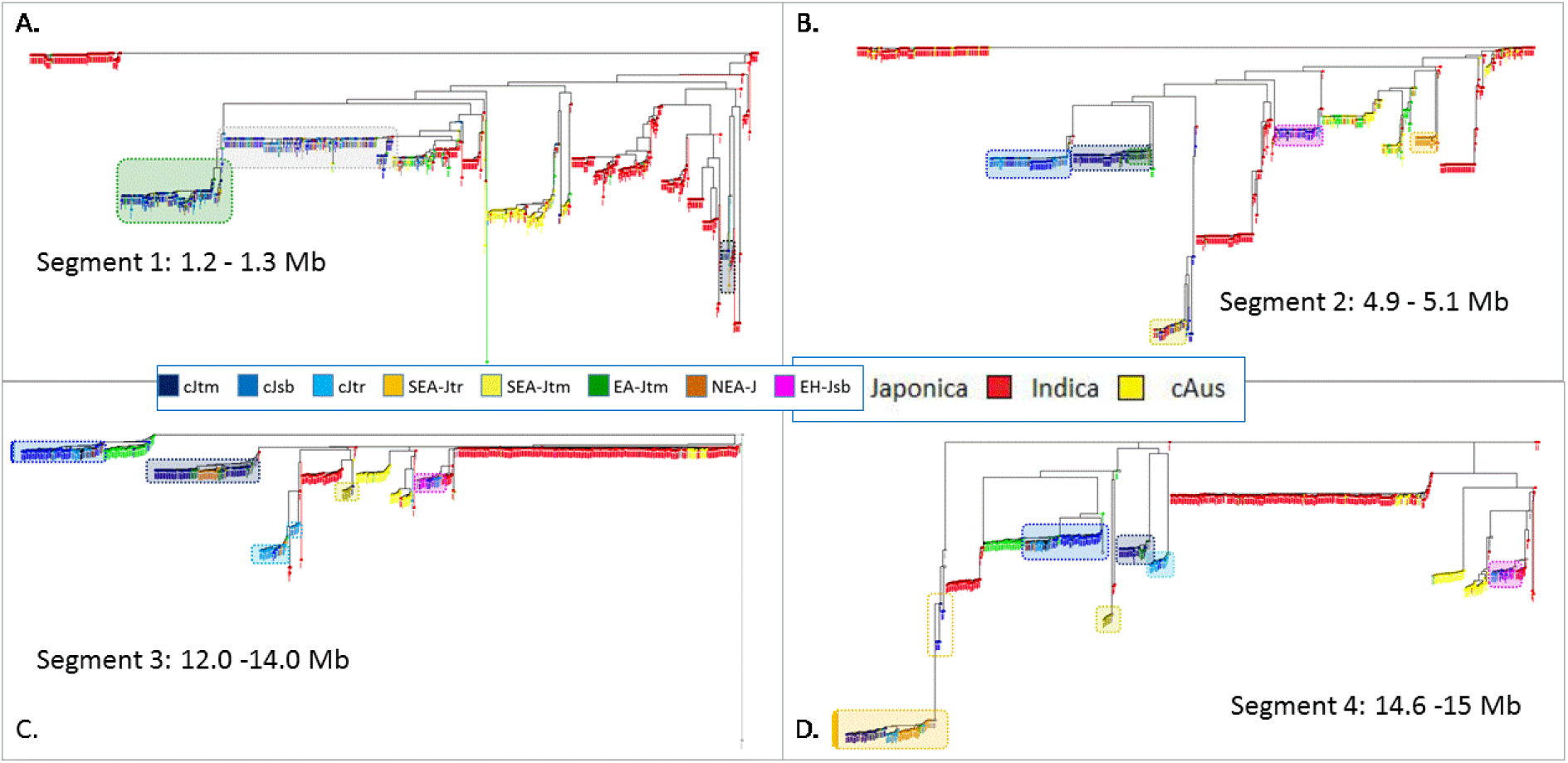
NJ trees of all CORE accessions derived from SNP analysis within targeted segments of chromosome 8. Segments were selected to reflect different streams detected across Japonica genomes through visual sampling of the admixture patterns in Fig. 5. Japonica accessions in NJ trees were attributed the MS subgroup colours in Fig. 3. Their local classification was assessed by visual inspection of ideogram patterns and synthesized by a coloured box reflecting this assignment. Indica and *c*Aus accessions are depicted in red and yellow, respectively.

Japonica varieties were distributed in five main branches when considering variation within the longest segment (12-14 Mb) (Fig. 5, 6C):

- one shared with *c*Basmati that clusters essentially common subtropical accessions;
- one consisting essentially of common temperate accessions together with the EA temperate type (green) and the SEA-Jtr tropical type (orange);
- one mostly consisting of common tropical types;
- one essentially consisting of the SEA temperate type, connected to an Indica branch;
- one that pools EH subtropical types close to an Indica branch of diverse origin and (Indica) subgroup classification.

Four smaller regions with broad distribution of specific segments on chromosome 8 illustrated the penetration of specific segments across germplasm compartments. When they are analysed in order of decreasing length, the following observations are apparent:

A 400 kb segment (14.6-15.0 Mb, Fig. 5, Fig. 6D) featured both a SEA-tropical (orange) segment and an EH-subtropical segment (pink) with a broad distribution far beyond their corresponding subgroup. Here also, both branches connected outside of the Japonica distribution (Fig. 6). EH-sb haplotypes were closely related to some Indica haplotypes, suggesting that this stream also extended to the Indica group. The remaining Japonica accessions fell essentially along three branches, roughly corresponding to the common temperate, subtropical and tropical types, yet with many individual exceptions, plus a fourth branch bearing haplotypes classified as SEA-Jtm (greenish yellow). The distance separating the diverse Japonica branches was as marked as that with *c*Aus or Indica domains. In this tree, *c*Basmati accessions fell within the Japonica section, close to the common subtropical accessions.

A shorter 200 kb segment (4.9-5.1 Mb, Fig. 5, Fig. 6B) featured a SEA-tropical (orange) segment well aligned to its corresponding subgroup and an EH-subtropical-marked segment (pink) with a broad distribution far beyond its corresponding subgroup. Indeed, the two corresponding branches were observed (Fig. 6). Both were rooted outside of the Japonica domain. A third noteworthy branch consisted of haplotypes classified as SEA-Jtm accessions together with a few other Japonica accessions and a few Indica accessions, suggesting that this distribution extended into the Indica group. This latter branch was markedly differentiated from the rest of the species. In this tree, *c*Basmati accessions were dispersed in the *c*Aus part of the tree.

The third case related to a 100 kb segment (1.2-1.3 Mb, Fig. 5, Fig. 6A) featuring a fine EA-Jtm signal (green) that was found in many tropical accessions. The specific NJ tree confirmed that these tropical accessions shared a common haplotype with the EA-Jtm accessions, which appeared at the edge of a long branch, far beyond the variation observed among the rest of the Japonica accessions (Fig. 6). The sequence data revealed concerted variation for 78 diagnostic SNP loci distributed between positions 1199565 (first) and 1291127 (last). Seven haplotype configurations were observed in relation to the green segments (Supplementary Table S2): the longest and most frequent one (91563 bp long), with the extremities as above, and six shorter ones, ranging from 87372 bp to 5867 bp in size. On the ideograms, the longest three (> 70 kb) appeared as green segments, while the other four (< 20 kb) appeared as very thin green lines. All these segments were concentrated among Japonica accessions (frequency of 44%) and were very rare in non-Japonica accessions (frequency <0.5%). The most widely represented segment fraction extended from 1203756 to 1206235 and spanned part of the PP2 gene, i.e. a heat-shock protein gene upregulated by drought (Jin *et al*. 2013). Another tree branch bearing fewer accessions was long, anchored outside of the Japonica group and corresponded to a haplotype identified as outlier in our former study. In this tree, *c*Basmati accessions clustered within the Japonica branch close to a small set of Indica accessions from Southeast Asia.

Many more examples of segments were investigated (see Fig. S8, S9). These patterns reflected variation localized at the end of branches, which were generally long and connected to an unstable upper level context. Overall, they highlighted marked differentiation from Indica, *c*Aus and common Japonica subgroups. The section below addresses this question of differentiation overall.

### Targeted distance analysis

Analysis of the distribution of the distances of selected clusters to the remaining domesticated diversity locally revealed a stable pattern across the subgroups considered. All five subgroups presented average differentiation above the local mean differentiation (Fig. S10). There was some variation, with the NEA tropical cluster presenting local Z scores averaging 0.715 and the SEA tropical cluster showed an average Z score of 1.052 (Fig. S10). The distributions of these distances were bell-shaped, albeit with heavy upper tails: from 17.2 % (NEA tropical) to 26% (for SEA tropical) of the clusters presented Z scores above 2 (Fig. S10). In Figure 7, the distances to CRG centroids are represented by the *y* axis. An analysis of the *x* axis revealed that distances to target classes accessions also hold elevated average scores.

**Fig. 7.**
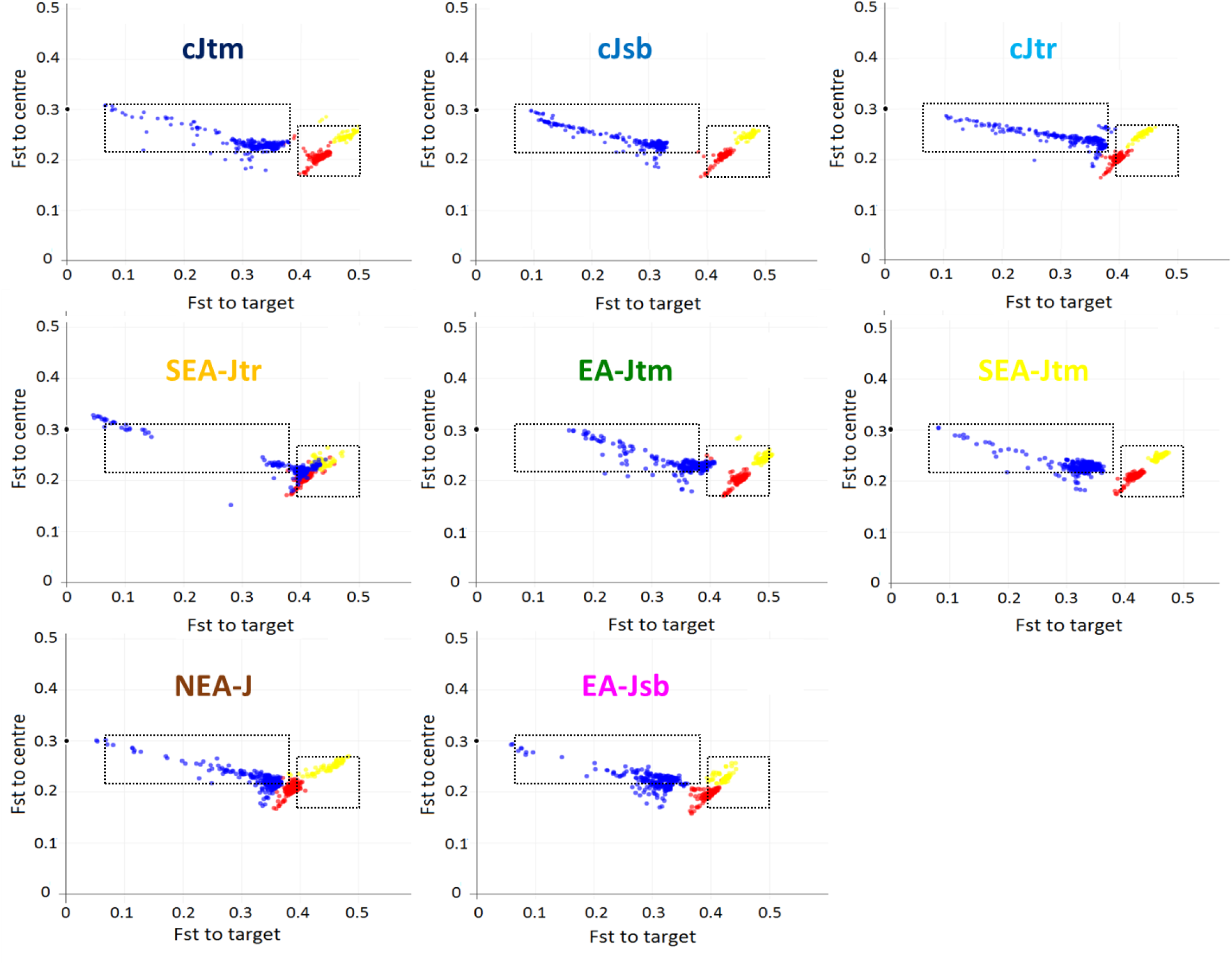
Bi-dimensional characterization of variation pertaining to different streams in Japonica germplasm. Class-specific target haplotypes were identified at each 150 SNP window through maximum affinity to the queried MS vectors and were used to derive a target centroid. The remaining haplotypes were used to derive a non-target CRG centroid. Distances to target and CRG centroids were calculated across windows in feature space and converted to *Fst* using a linear regression in logarithmic space. Distances to the target centroid (*x* axis) were plotted against distances to the proxy CRG centroid (*y* axis). The figure shows the distribution of CORE reference accessions using the various streams sequentially. The colour codes for scatter plots are as in Figure 2A. The colour codes for class designations are as in Figure 3. Empty boxes were included to facilitate comparison among streams.

The distributions of accessions in Fig.7 enable inferences in terms of introgression from a given source: 1) a steep negative slope suggests a highly divergent source; 2) a clear separation of the Japonica bulk from the Indica and *c*Aus bulks suggests a source closer to the former; 3) a Japonica range reaching low distances to the target suggests a high concentration of this source in some accessions; and conversely, 4) a discontinuity in the Japonica distribution indicates significant isolation of the target subgroup.

These interpretations suggest that the EA-Jtm stream is related to a highly divergent source that is currently more dissipated than the others. The SEA-Jtr stream also reveals a highly divergent source which, moreover, is as divergent from Japonica as it is from the others, and which seems concentrated in a markedly isolated subgroup. The NEA-J stream also suggests a source external to Japonica, while the EH-Jsb stream appears less divergent but more confined. cJtr and cJsb distributions are characterized by a slighter slope and cJtr also by a very limited offset of the bulk of Japonica vs Indica/*c*Aus. Based on the assumption that introgression is usually followed by assimilation, i.e. dilution of the corresponding source contribution and (re-)establishment of an equilibrium state, the three SEA-Jtr, NEA-J and EH-Jsb components would be of more recent origin than the others and EA-Jtm would be the most ancient. It is among the first that we found those that jointly differentiate from Indica, *c*Aus and Japonica (SEA-Jtr and NEA-J).

### Japonica component of cBasmati varieties

Considerable proportions of *c*Basmati genomes were found to be specifically related to Japonica variation, as evidenced by the summary of local window classifications (Table 1): mean= 28 %, min = 15.5 and max = 34.2. We looked at how these regions were distributed among Japonica specific classes and found that significant portions of *c*Basmati genomes were discriminant of Japonica classes, starting with cJtm, cJsb and cJtr (median: 27.7 Mb, 21 Mb and 7.6 Mb), while also including the other classes in close proportion to those observed for them in Japonica (Figure 8).

**Fig. 8.**
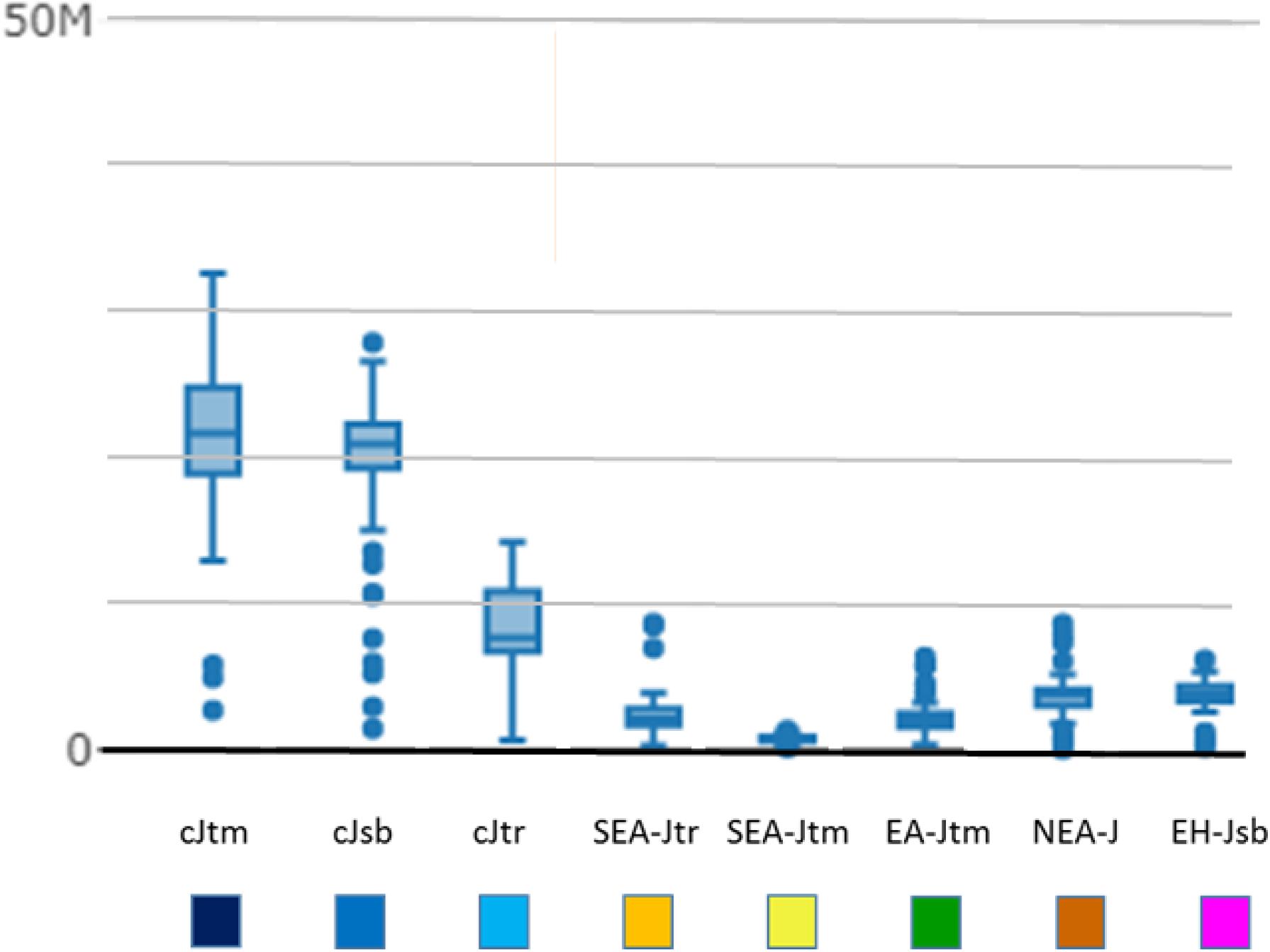
Distribution of *c*Basmati genome segments assigned to the various Japonica classes. Haplotype size (in bp) was summed by class and individual accession and analysed across *c*Basmati varieties and Japonica classes.

While temperate Japonica haplotypes appeared to be uniformly distributed across chromosomes, tropical and subtropical haplotypes showed uneven distributions, with most contributions concentrated on chromosomes 4, 8 and 9 and chromosome 8, respectively (Supplementary Figure S8). Consistent signatures pertaining to Japonica sub-differentiation were also noted. All sub-groups were represented among *c*Basmati haplotypes to some extent and covering, except for SEA-Jtm, medians above two megabases of the respective genomes (lower quantiles above one Mb with the same exception). Although the distribution was not fully uniform, most sources appeared in significant length along most chromosomes (Supplementary Figure S7).

In addition to these summary statistics, Figure 5 shows how the central portion of chromosome 8 in *c*Basmati displayed a physical structure among specific haplotypes that combined at least three Japonica vertices with a unique massive configuration. On the same chromosome, cases where *c*Basmati displayed a Japonica-affine segment (segments 3 and 4 in Figure 6), those borne by *c*Basmati accessions and the other Japonica versions had diverged enough for the *c*Basmati accessions to form a distinct cluster.

## Discussion

We explored genetic variation patterns within a group of Asian cultivated landraces representative of Japonica, one of the major and possibly oldest, domesticated rice groups. Japonica was expected to be the simplest rice group, derived directly from a domestication event in China and characterized by lower diversity than all other groups.

The classification history within this group has always had a geographical component. Initial classification schemes recognised temperate and tropical forms differentiated essentially on the basis of morphological traits (Morinaga 1954, Oka 1958), hybrid fertility relationships with Indica (Morinaga and Kuriyama 1958) and molecular marker findings (Garris *et al*. 2005). Some studies went as far as identifying a “Javanica” type at the same level as Japonica and Indica (Matsuo 1952, Morinaga and Kuriyama 1958), while others placed Javanica within a large Japonica group and highlighted the importance of samples from South and Southeast Asia, which often appear to be intermediate in terms of morphology (Glaszmann and Arraudeau, 1986) while being rich in specific alleles at marker loci (Glaszmann 1988). The use of larger sampling sizes and immensely larger numbers of molecular polymorphisms led to the recognition of a single Japonica group including temperate and tropical subgroups, and a third subgroup encapsulating subtropical variation from Southeast Asia (Wang *et al*. 2018).

In order to gain greater insight into this genepool, we relied on an initial classification of local genomic haplotypes in reference groups (Santos *et al*. 2019) so as to focus solely on genome regions free of recent introgressions from Indica and *c*Aus. We used PCA-based kernel density estimates for haplotype classification along the genome in order to select only derived variables bearing maximum affinity with Japonica. After testing the capacity of a random selection of PCA-derived variables to successfully retrieve the main foci of global rice diversity, we showed that profiles selected for Japonica classification were able to retrieve known diversity components and conducted an in-depth study on the structure within this target group.

We thus highlight genomic patterns characteristic of the first components of the Japonica variation, i.e. temperate, tropical and subtropical layers on the first PCA plane, as well as those appearing on subsequent axes. The third axis clearly isolated a subgroup of tropical accessions from Indonesia representing about 10% of the whole Japonica sample size. The position of Japonica subgroups in this analysis mimics the historical progression of their discovery, with temperate and tropical Japonica explained on the first axis, subtropical on the second, and Indonesian tropical, as emerged in a more recent clustering conducted based on the 3K-RG dataset (Supplementary Figure S11), on the fourth axis. Subsequent axes reveal finer components whose visible representatives are few in number and have localised geographical distributions. Some of these components are concentrated in a single of the major subgroups, others transcend this classification.

We characterised the variation borne by these components relative to core reference accessions, including Japonica, Indica and *c*Aus accessions. All eight components show an average distance to core centroids exceeding local differentiation among other domesticates, indicating that the surveyed clusters hold peripheral positions within the global rice structure. Some seem to be markedly more differentiated than others or more confined to specific varietal origins, while others appear to be more dissipated throughout the germplasm. We find evidence of true introgression from differentiated sources in the form of physically clustered signals conserved among focus accessions. While common temperate, subtropical and tropical variation shows specific patterns that are roughly dispersed over the whole genome, those components that appear in subsequent steps are increasingly localized, appearing more as streams of introgression in the global genomic topography rather than coherent structural entities. The simplest dependency sequence that could explain this pattern involves the derivation of the common subtropical subgroup from the temperate subgroup through the incorporation of an additional source, followed by the emergence of the tropical subgroup augmented by another additional source. Taken together, the genetic distance and genomic distribution variables thus make a strong case in favour of independent differentiation of the material considered, with subsequent incorporation into the Japonica genepool.

The current geographic distribution of these signals provides further evidence of profuse movements of rice varieties in many directions. Yet, it is not clear whether the higher signal concentration in a single genome could be indicative of closer proximity with the actual origin of the introgression event. Detailed case-by-case introgression studies would now be required to build a meaningful global picture.

Figure 9 presents a schematic representation of Japonica diversity as perceived on the basis of the findings of our study.

**Figure 9.**
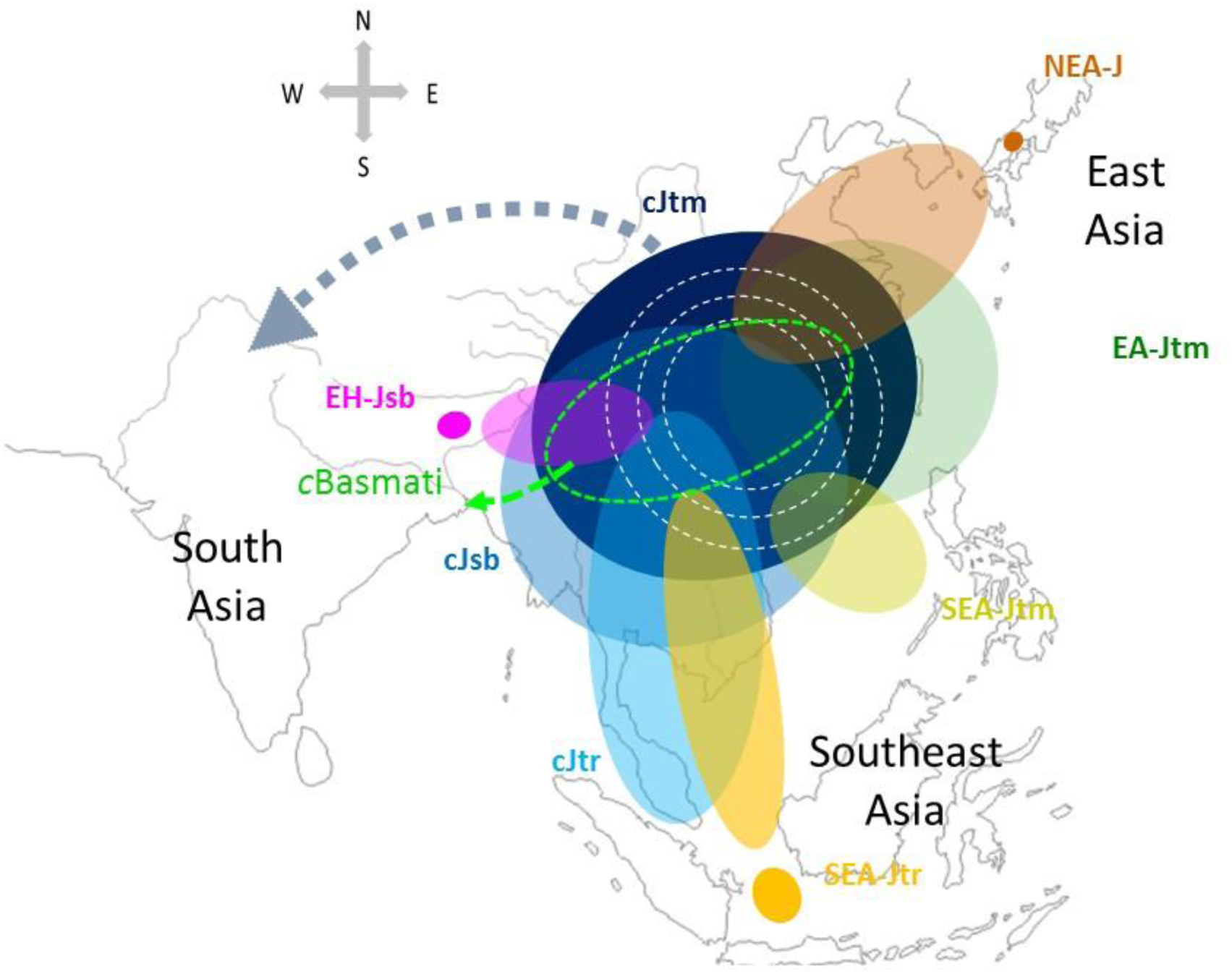
Schematic representation of genomic streams within Japonica rice. From a central origin in China, Japonica was augmented with a sequence of incorporations leading to three major groups (cJtm, cJsb and cJtr), as well as introgressions at the periphery of its distribution (EA-Jtm, SEA-Jtm, NEA-J,SEA-Jtr and EH-Jsb). The most recent ones are marked by clearcut subgroups (coloured disks). Profuse movements (concentric white circles) ensure coherence through exchanges between the various forms. The blue arrow designates migration from China to South and West Asia via the Silk Road. The light green arrow designates the contribution of Japonica to *c*Basmati. Map background from Daniel Dalet.

By the end of the second millennium BC domestication traits are evident north of the Yangtze River, in the Sichuan Basin and Yunnan, and south, in the Liaoning Peninsula region and Korea (Fuller *et al*. 2010). Our findings of differentiated material obtained by combining sources as diverse as EA-Jtm, SEA-Jtr and EH-Jsb indicate the integration of local rice variation into an expanding domesticated genepool. Thus, this appears to have been a plausible time period during which this material was first integrated into the domesticated genepool, as further supported by archaeological evidence of the actual timing of this expansion.

*c*Basmati is a commercially important group of rice varieties originating from the Himalayan foothills region and long proposed to be the result of contacts between major rice groups and local differentiated material (Glaszmann 1987, Garris *et al*. 2005). Fuller *et al.* (2010) placed the timing of these contacts in the first century AD, concomitant with a westward push, from the Ganges region, of admixed rice varieties bearing Japonica, *c*Aus, and possibly even Indica contributions (Jain *et al.* 2004). In a previous study, we specifically assessed variations in *c*Basmati varieties in relation to Japonica, *c*Aus and Indica genepools (Santos *et al*. 2019). At the same time, Civáň *et al*. (2019) inferred that the admixture occurred 4,000– 2,400 years ago, soon after Japonica rice reached the region (Bates *et al.* 2017). Today, the location of haplotypes significantly discriminant of Japonica across *c*Basmati genomes has been well characterised. The part of the genome putatively derived from Japonica in the hybrid make-up of *c*Basmati varieties represents a specific example of the Japonica diversity that was isolated at the time of the initial hybridizations. We found evidence of contributions from all major vertices of the Japonica substructure within this group, including evidence of cryptic variations specific to Indonesian, Japanese and Korean varieties. This combination of signals from geographically distant sources within the Japonica group, together with linkage patterns conserved among the *c*Basmati, suggests that this contribution occurred after the aggregation of all sources of diversity in Japonica subspecies. Given the upper bound inferred by Civáň *et al*. (2019), and the timing predictions of Fuller *et al.* (2010), current evidence thus seems to suggest that circum-Basmati varieties emerged sometime during or shortly after the last millennium BC. This is an important date as it provides an upper bound regarding when the Japonica genepool could have become internally connected.

## Conclusion

By comparing the findings of small-scale analyses conducted along the genome, we gained access to a fine-grained description of genetic entities previously described only through their genome-wide correlations. In pursuing this avenue of research, we have expanded current knowledge on rice genomic diversity by revealing several new pockets of diversity within the large well-known Japonica cluster. We found evidence of the incorporation into the Japonica genepool of cryptic variation of alien origin that is currently concentrated in several small subgroups of varieties scattered throughout Asia, and we traced this variation throughout the genome of the Japonica genepool. This newly uncovered variation represents 5–20 % of genome regions specific to Japonica. By extending our analysis to cover *c*Basmati genomes, we found evidence indicating that contacts between these sources pre-date the formation of this group. Our description paves the way for a more complete and systematic description of domesticated rice genetic variation, thus enhancing insight into its history. The analysis could be repeated using, for read mapping, reference genomes other than that of Nipponbare to help bridge the gap between any single reference genome and the rice pangenome (Zhao et al 2018, Wang et al 2018). It is indeed plausible that many signals useful for reconstructing the streams of introgression accessible through the 3K-RG dataset are scattered among reads that are unmapped when Nipponbare is used as the sole reference. Our analysis could also be tailored for other groups, with a priority to the globally very diverse Indica group. It could also be further developed to take advantage of genome regions that are shared by two groups or more. Application of our strategy to other crops could also likely reveal equally complex scenarios and open new avenues for documenting crop evolution and its coevolution with humankind. Our conclusions on Japonica rice stress the importance of constantly broadening the genetic base of crops to enhance their adaptation.

## Supplementary Material

**Supplementary Table S1** - Group assignment of Japonica accessions using MS group profiles.

**Supplementary Figure S1** – PCA Euclidian distances and genetic correlation.

**Supplementary Figure S2** – PCA and K-means analysis of MAC variables selected for Japonica classification, PCs 1-3.

**Supplementary Figure S3** – PCA and K-means analysis of MAC variables selected for Japonica classification, PCs 3-5.

**Supplementary Figure S4 –** PCA and K-means analysis of MAC variables selected for Japonica classification, PCs 6-10.

**Supplementary Figure S5 –** MAC assignment to groups indicative of Japonica substructure along the genome of tropical Japonica accessions.

**Supplementary Figure S6** – MAC assignment to groups indicative of Japonica substructure along the genome of tropical Japonica accessions.

**Supplementary Figure S7** – MAC assignment to groups indicative of Japonica substructure along the genome of tropical admixed accessions.

**Supplementary Figure S8** – Local genomic imprints of Japonica sub-taxa for chromosomes 1 and 10.

**Supplementary Figure S9 –** Local genomic imprints of tropical Japonica sub-taxa along chromosome 6.

**Supplementary Figure S10** – Distributions of normalized distances to CRG bodies of variation.

**Supplementary Figure S11** – MDS plot of Japonica lines (n=854) based on 3K filtered SNP set.

## Data Availability

All the code used in this study is available online (DOI: 10.5281/zenodo.3142884).

## Acknowledgements

This study was funded by CIRAD with support of the project Genome Harvest, reference ID 1504-006, through the “Investissements d’Avenir” program (Labex Agro: ANR-10-LABX-0001-01), the project AdaptGrass, reference ID170544IA, through the “Investissements d’Avenir” program (I-SITE MUSE: ANR-16-IDEX-0006) and the CGIAR Research Program on Rice Agrifood Systems (RICE). This work was supported by the CIRAD - UMR AGAP HPC Data Center of the South Green Bioinformatics platform (http://www.south-green.fr/).

