## Supplementary Figure S8 for "Multiple streams of genetic diversity in Japonica rice"

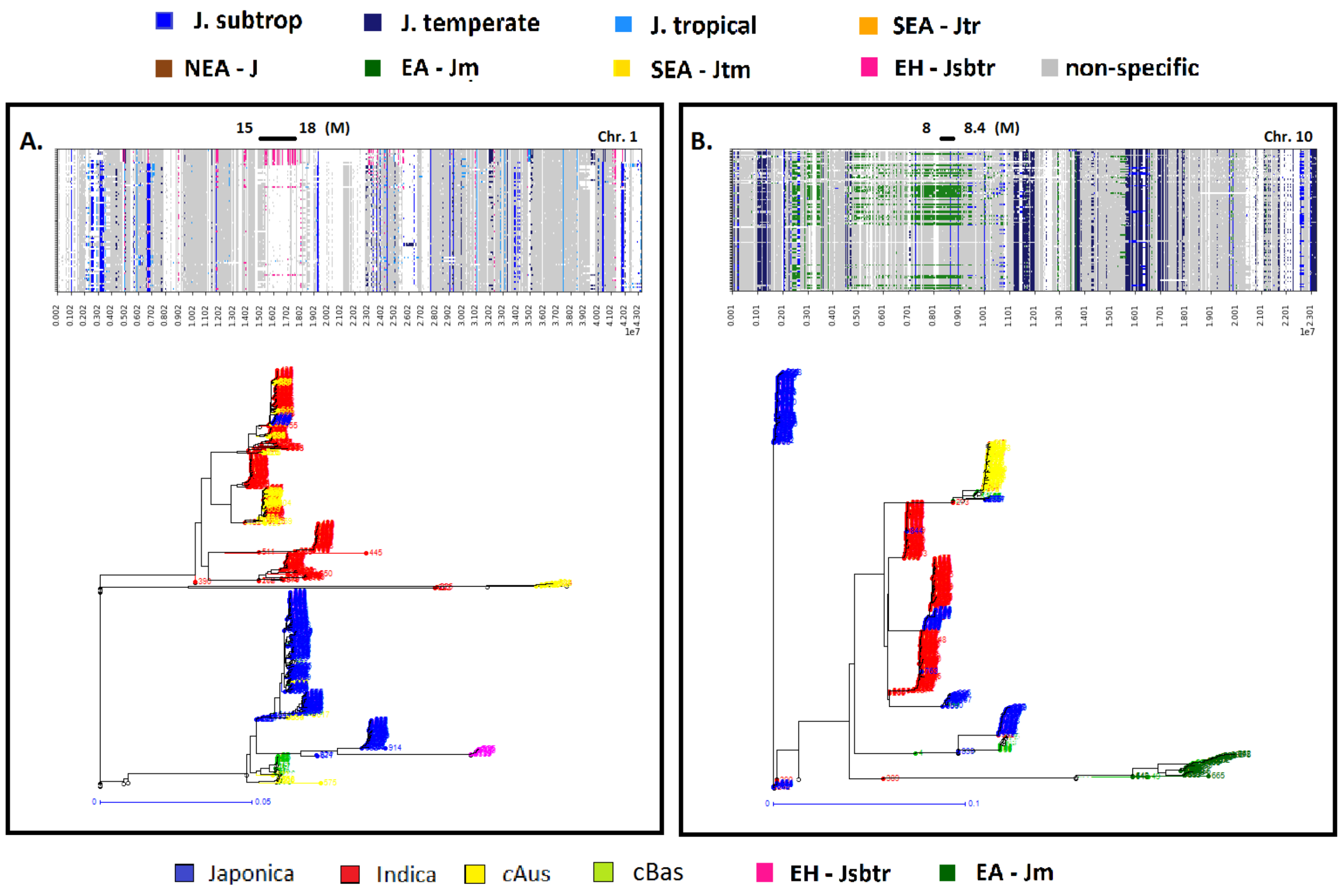


**Fig. S8 Local genomic imprints of Japonica sub-taxa for chromosomes 1 and 10.** Average profile of vector groups was used to select for correlation for Japonica sub-taxa of subtropical Japonica accessions from Bhutan and temperate Japonica accessions mostly from China. MAC assignment to vectors in groups identified is plotted in ideogram format. Local regions were selected visually. SNP data was extracted and used to construct neighbour-joining trees, plotted bellow the respective ideograms. Individuals were first labelled according to their global group of assignment (Japonica, Indica, *c*Aus). Accessions bearing sub-taxa specific local haplotypes within the regions extracted were then labelled according to their local labels. **A**) Sub-taxon signals across subtropical Japonica chromosome 1. NJ-trees of SNP data extracted between Mb 15 and Mb 18 **B**) Sub-taxon signals across tropical Japonica chromosome 10. NJ-trees of SNP data extracted between Mb 8 and Mb 8.4. Accessions in NJ trees were attributed MS group colour code by visual inspection of ideogram patterns across Core accessions.
