## Supplementary Figure S9 for "Multiple streams of genetic diversity in Japonica rice"

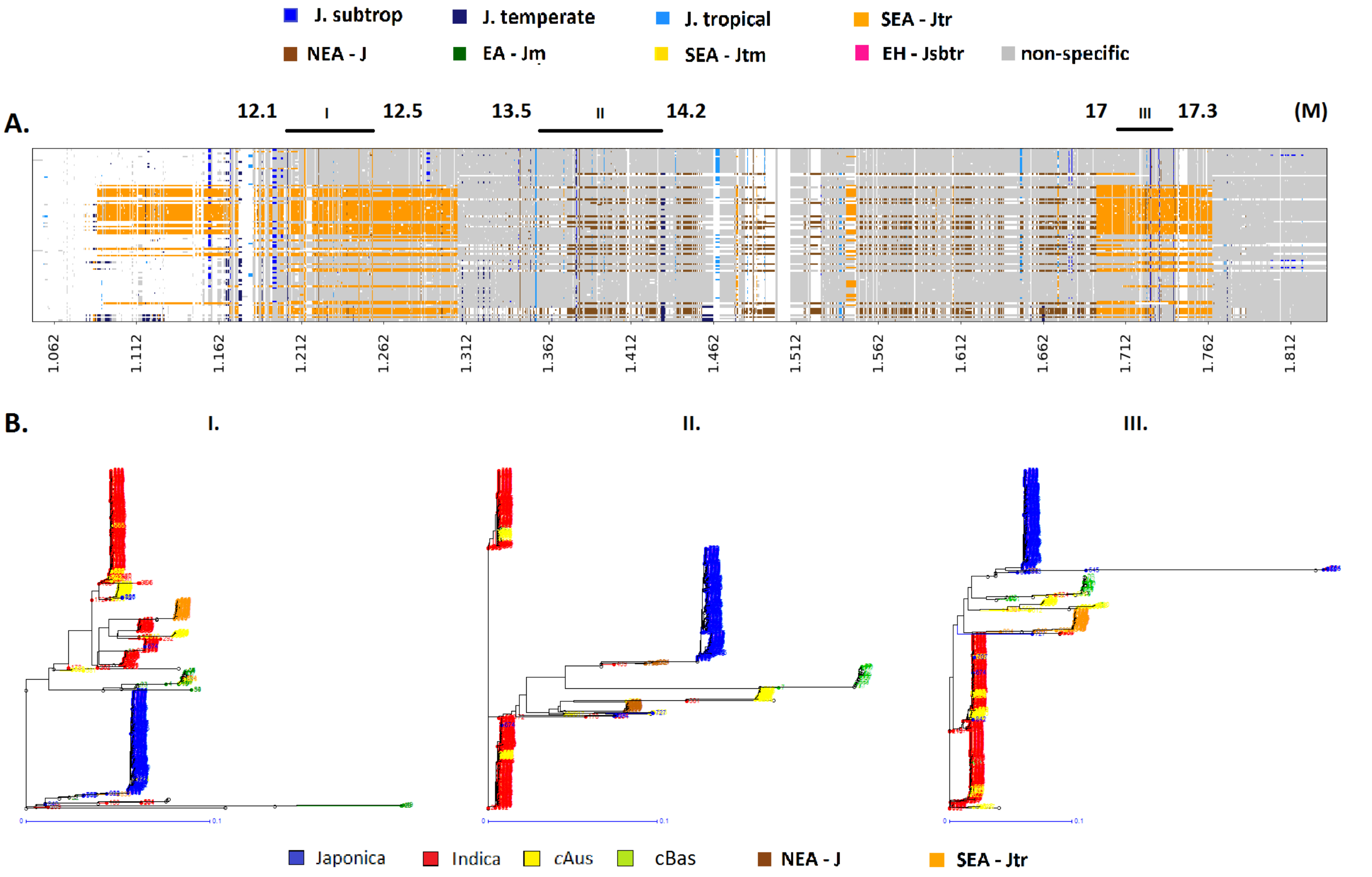


**Fig. S9 Local genomic imprints of tropical Japonica sub-taxa along chromosome 6.** Average profile of vector groups was used to select for correlation for Japonica sub-taxa of subtropical Japonica accessions from Bhutan and temperate Japonica accessions mostly from China. MAC assignment to vectors in groups identified is plotted in ideogram format. Local regions were selected visually. SNP data was extracted and used to construct neighbour-joining trees, plotted bellow the respective ideograms. **A**) Sub-taxon signals across tropical Japonica chromosome 6. Regions selected for extraction indicated above ideogram in millions of base pairs. **B**) NJ-trees of SNP data extracted at regions indicated in A. Individuals were first labelled according to their global group of assignment (Japonica, Indica, *c*Aus). Accessions in NJ trees were attributed MS group colour code by visual inspection of ideogram patterns across Core accessions.
