## Supplementary Figure S10 for "Multiple streams of genetic diversity in Japonica rice"

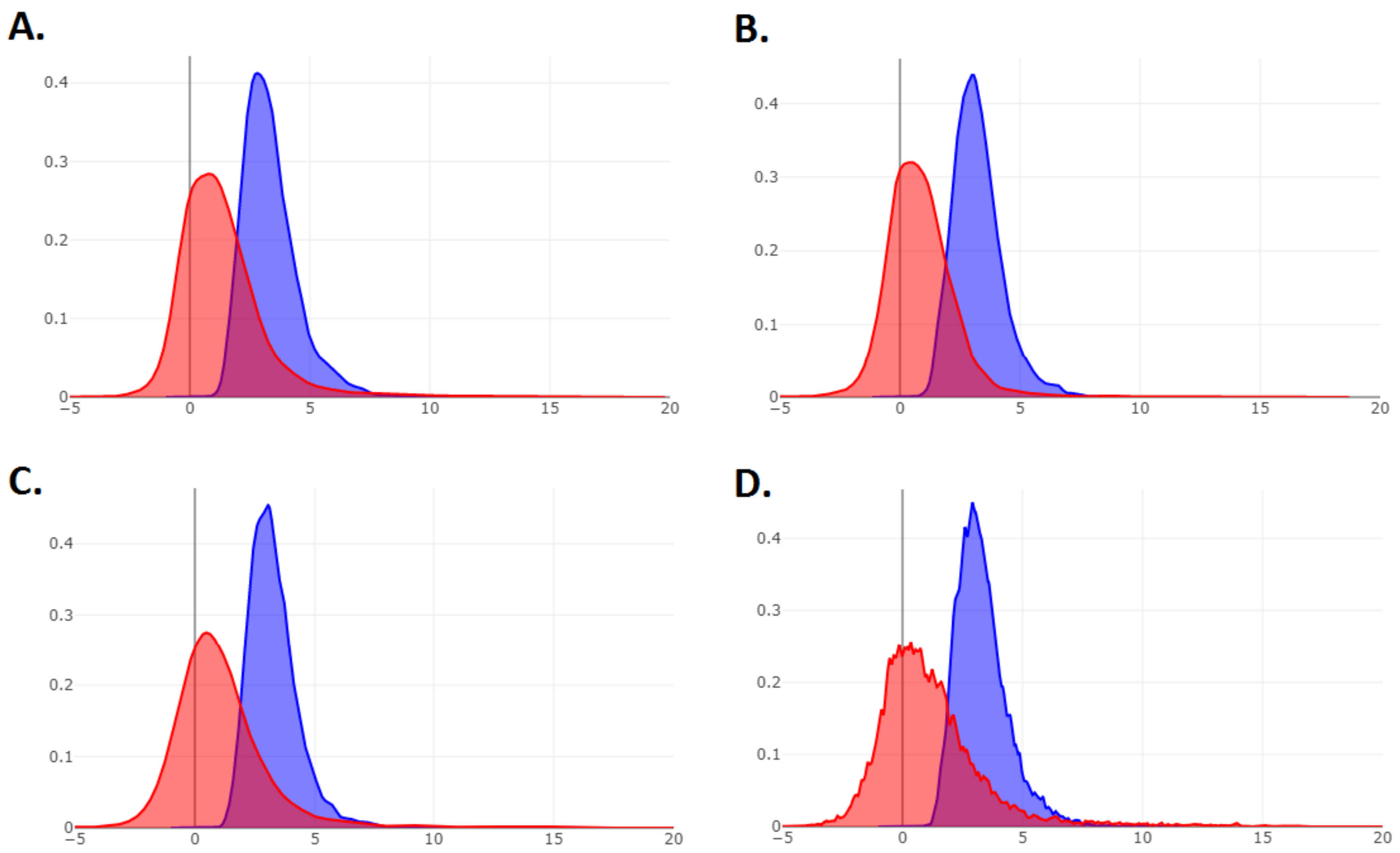


**Fig. 10 Distributions of normalized distances to CRG bodies of variation**. Distances to CRG and target haplotype centroids were calculated across windows in feature space. All measurements were standardized locally using the distance of non-target CRG accessions to proxy reference centroid. Normalized averages of local target distances to proxy CRG centroids are plotted in red. The blue curves depict the distribution of raw average distances to the same centroid among non-target CRG accessions. **A**) Distance distributions for clusters of SEA tropical Japonica accessions (Fig. 3B). **B**) Distance distributions for clusters of NEA tropical Japonica accessions (Fig. 3D). **C**) Distance distributions for clusters of EH Himalayan subtropical Japonica accessions (Fig. 3E). **D**) Distance distributions for clusters of EA temperate Japonica (Fig. 3C).
