## Supplementary Figure S11 for "Multiple streams of genetic diversity in Japonica rice"

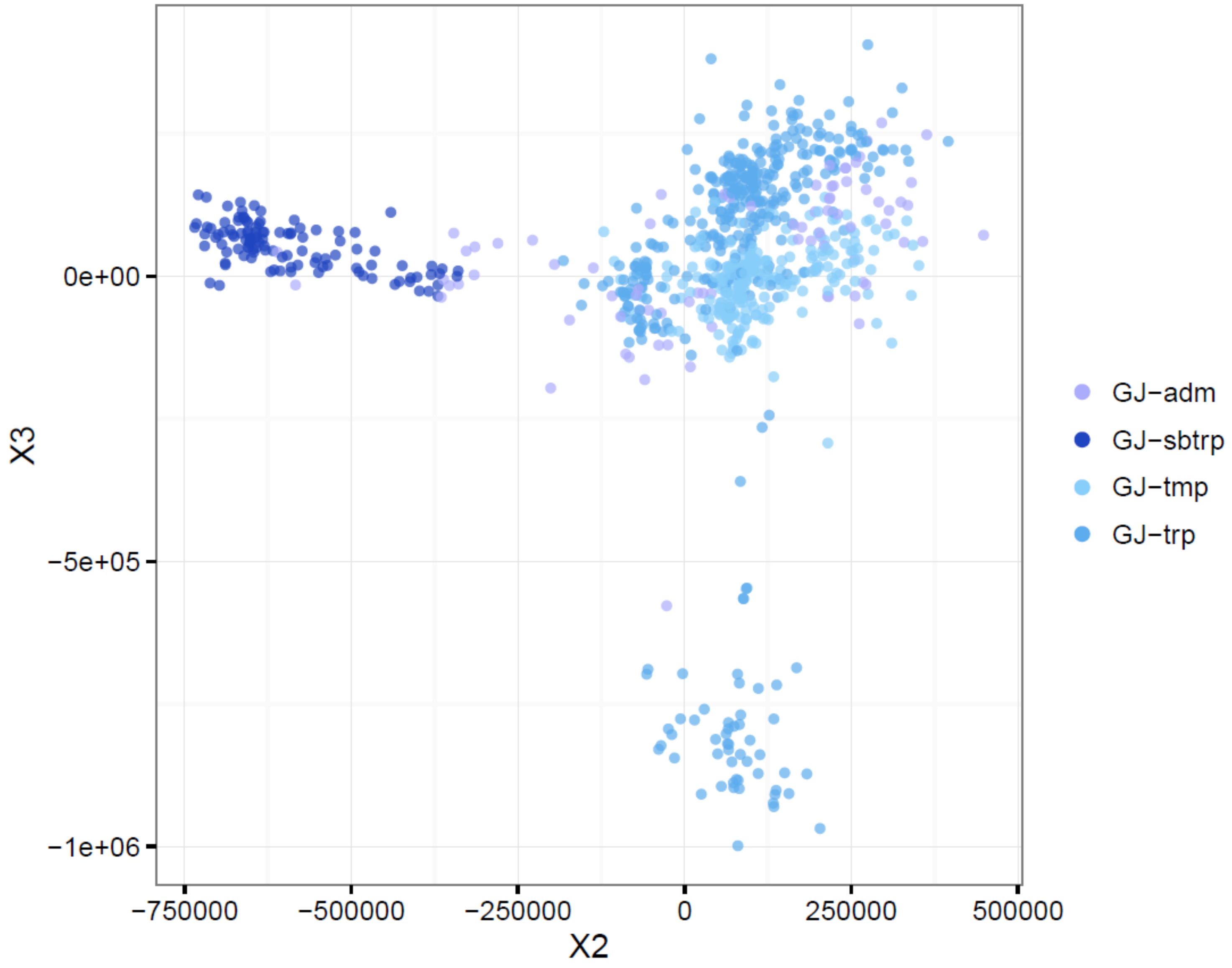


**Figure S11. MDS plot of Japonica lines (n=854) based on 3K filtered SNP set.** Multidimensional scaling was run on IBS distance matrix normalised into proportions instead of allele count. The colours denote admixture-based grouping from Wang *et al*. 2018: temp: temperate japonica; trop: tropical japonica; subtrop: subtropical japonica; japx: within-japonica admixed varieties.
