## Supplementary Figure S1 for "Multiple streams of genetic diversity in Japonica rice"

### Supplementary Figures

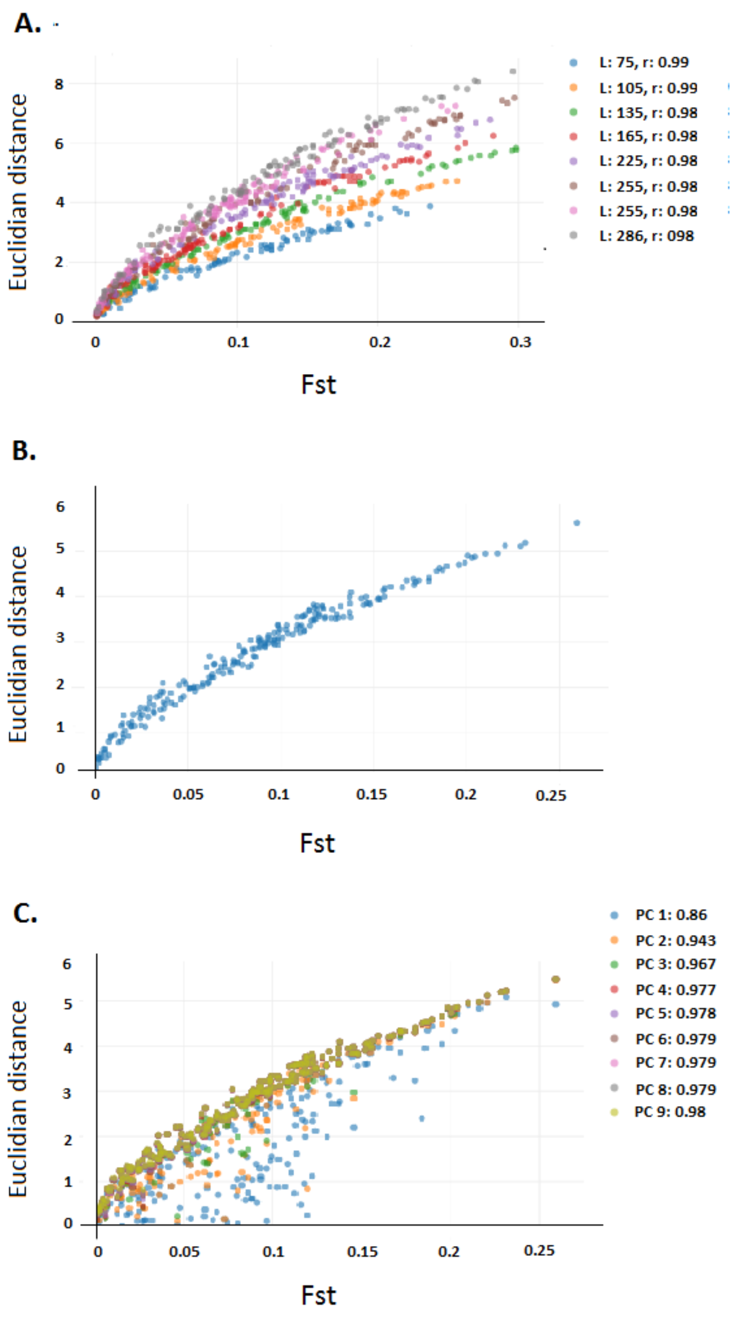


**Fig. S1. PCA Euclidian to Feature Space Distances.** Groups of eight vectors were selected at random from the base frequency data set. Genetic distances were calculated between pairs of vectors in $F_{st}$s. Haploid populations were sampled from each vector. Sampling was a random draw between 50 and 300. PCA was performed on haplotypes from all populations combined. Euclidian distances were calculated between the centroids of observations from all pairs of vectors using an incrementing number of components (1 to 9). **A)** Genetic distances *vs.* Euclidian distances in five-dimensional feature space for vectors of length 75 to 285 at steps of 20. **B)** Genetic distances *vs.* Euclidian distances in five-dimensional feature space for vectors of length 150. **C)** Genetic distances against their Euclidian equivalents calculated using one to 9 dimensions for vectors of length 150.
