## Supplementary Figure S2 for "Multiple streams of genetic diversity in Japonica rice"

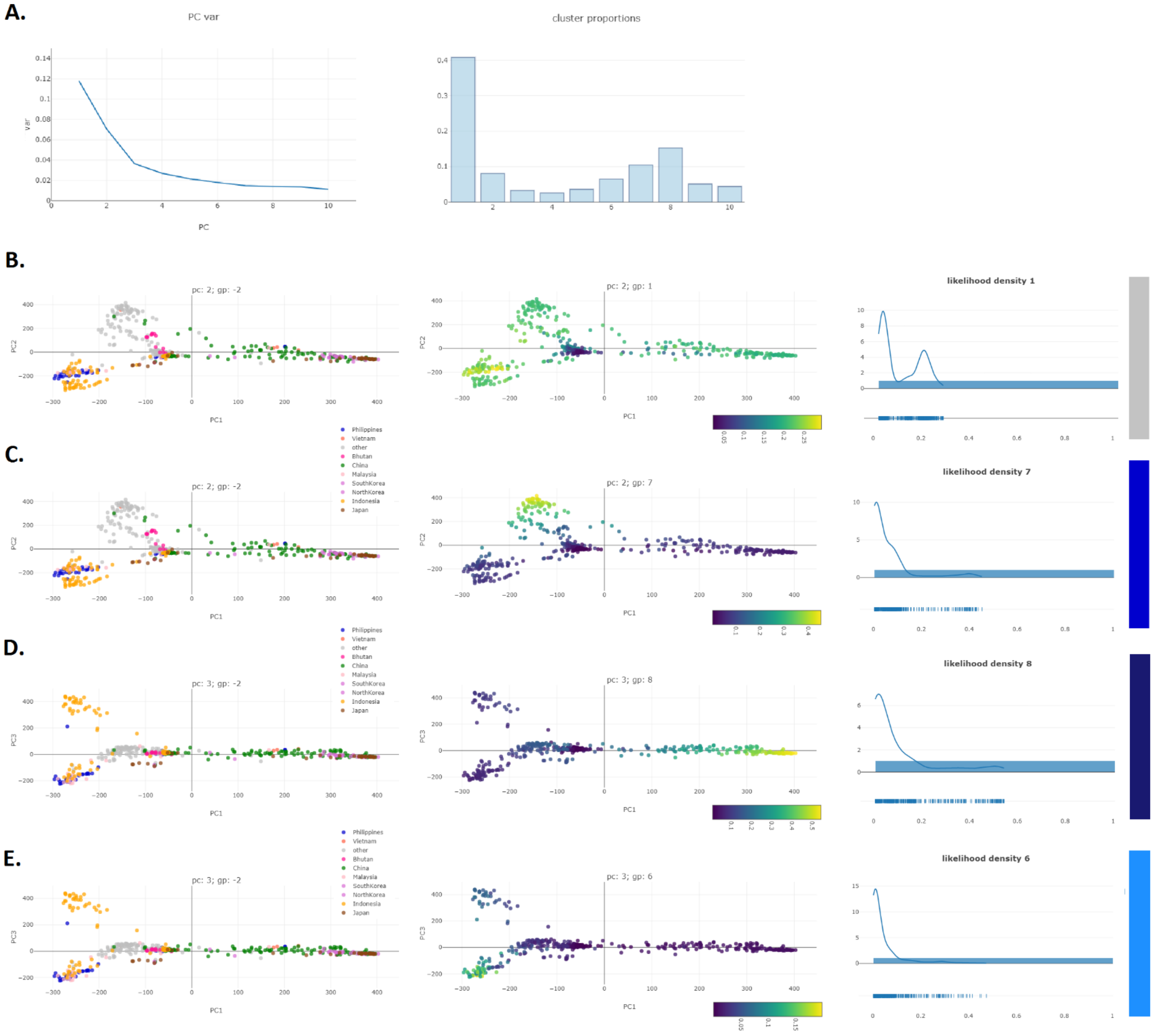


**Fig. S2 PCA and K-means analysis of MAC variables selected for Japonica classification, PCs 1-3.** Mean shift (MS) profiles of local clusters were selected for Japonica classification of maximum affine individual haplotypes. PCA was conducted on matrix of Japonica accessions x MS variables. Remaining Core accessions were projected using the resulting transformation. MS vector loadings were grouped using K-means at K = 10. Accession-level p-values were averaged by k-means group. **A**) Proportion of variance explained by principal component (left) and proportion of MS vectors assigned to each k-means group (right). **B**) Principal components 1 & 2 Core projections; individual average p-value across k-means group I; density plot of average p-values for group I. **C**) Principal components 1 & 2 Core projections; individual average *p*-value across k-means group VII; density plot of average p-values for group VII. **D**) Principal components 1 & 3 Core projections; individual average *p*-value across k-means group VIII; density plot of average *p*-values for group VIII. **E**) Principal components 1 & 3 Core projections; individual average *p*-value across k-means group VI; density plot of average *p*-values for group VI.
