## Supplementary Figure S3 for "Multiple streams of genetic diversity in Japonica rice"

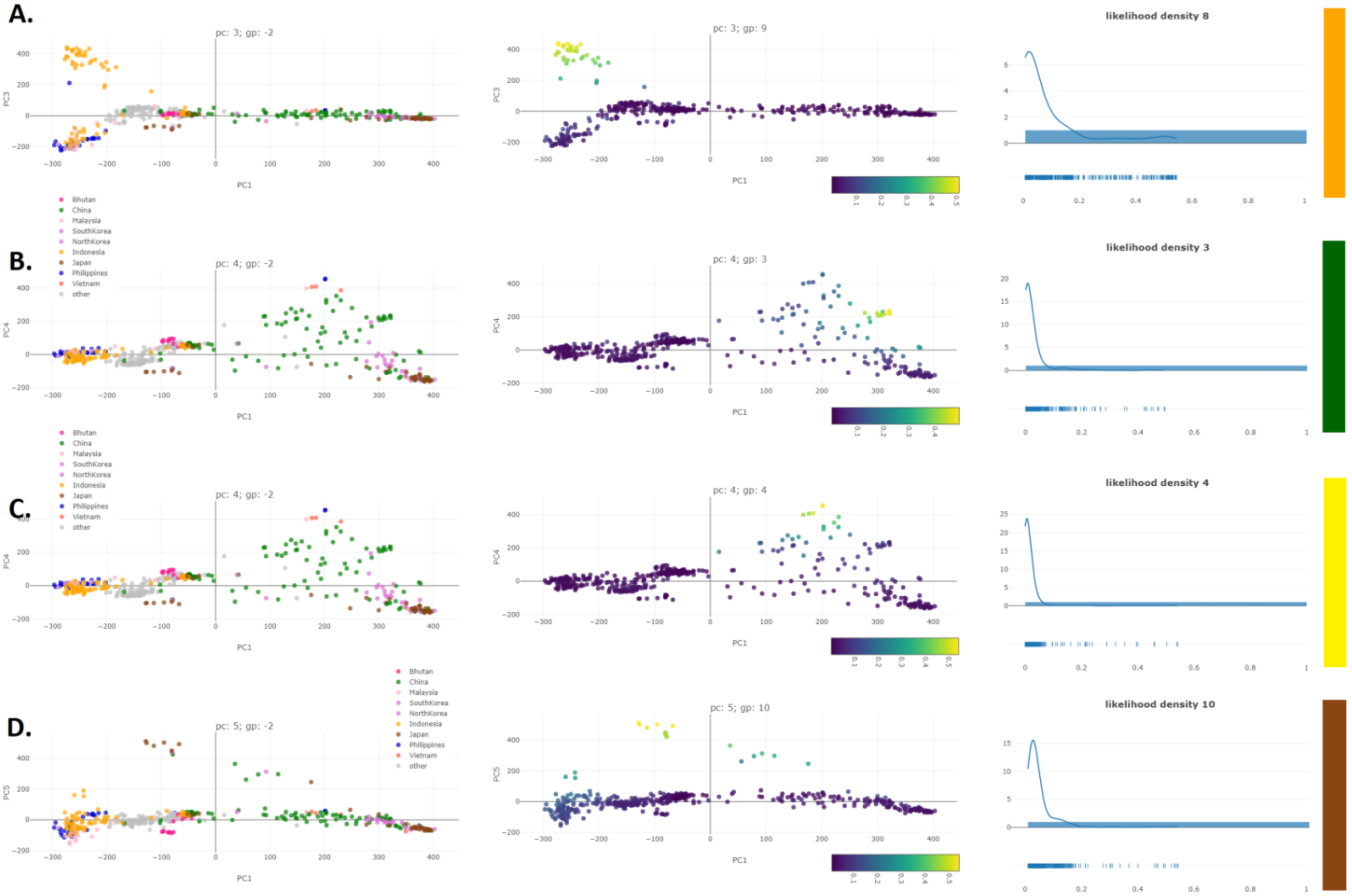


**Fig. S3 PCA and K-means analysis of MAC variables selected for Japonica classification, PCs 3-5.** Mean shift (MS) profiles of local clusters were selected for Japonica classification of maximum affine individual haplotypes. PCA was conducted on matrix of Japonica accessions x MS variables. Remaining Core accessions were projected using the resulting transformation. MS vector loadings were grouped using K-means at K = 10. Accession-level p-values were averaged by k-means group. **A**) Principal components 1 & 3 Core projections; individual average p-value across k-means group IX; density plot of average p-values for group IX. **B**) Principal components 1 & 4 Core projections; individual average *p*-value across k-means group III; density plot of average p-values for group III. **C**) Principal components 1 & 4 Core projections; individual average *p*-value across k-means group IV; density plot of average *p*-values for group IV. **D**) Principal components 1 & 5 Core projections; individual average *p*-value across k-means group X; density plot of average *p*-values for group X.
