## Supplementary Figure S4 for "Multiple streams of genetic diversity in Japonica rice"

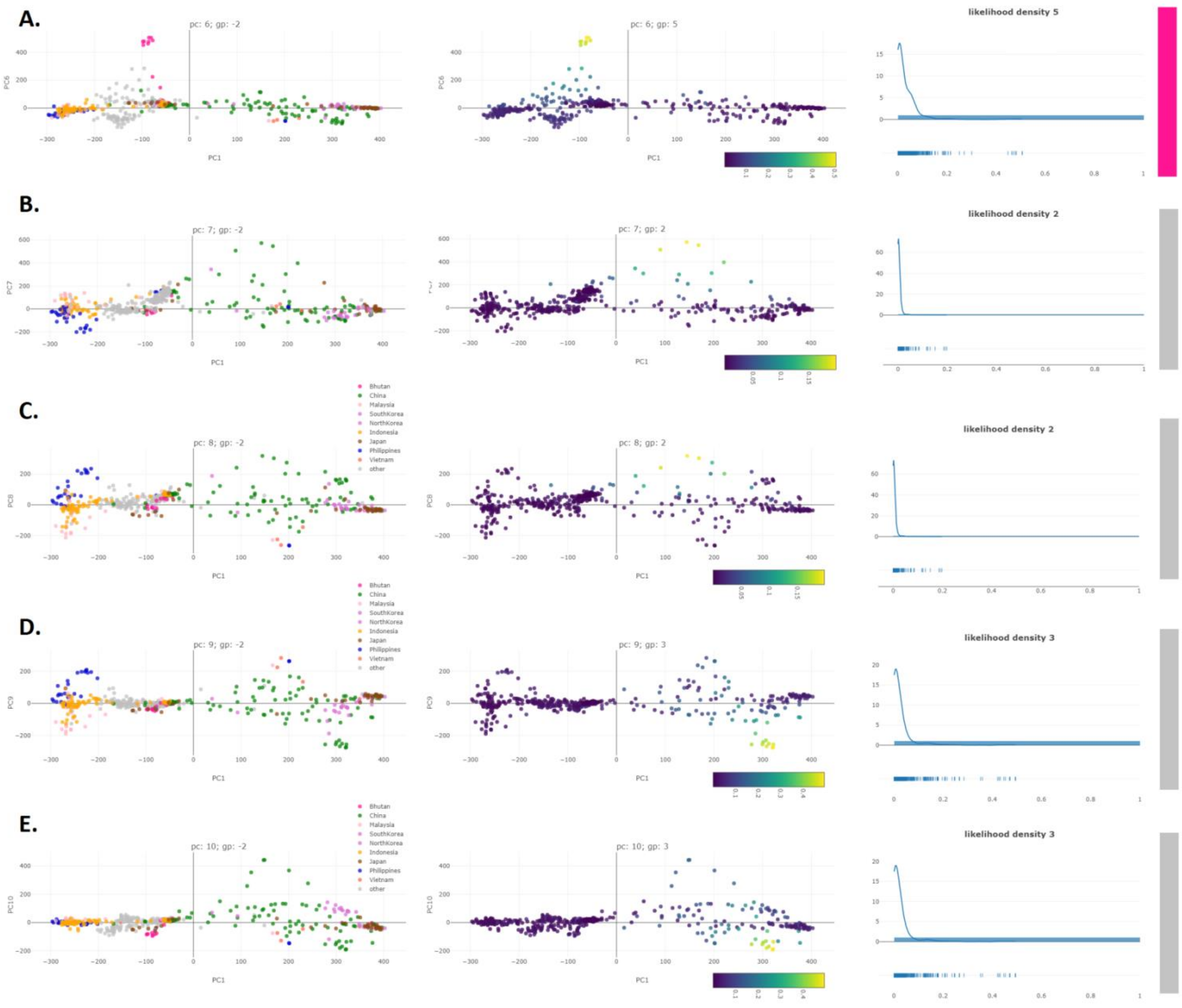


**Fig. S4 PCA and K-means analysis of MAC variables selected for Japonica classification, PCs 6-10.** Mean shift (MS) profiles of local clusters were selected for Japonica classification of maximum affine individual haplotypes. PCA was conducted on matrix of Japonica accessions x MS variables. Remaining Core accessions were projected using the resulting transformation. MS vector loadings were grouped using K-means at K = 10. Accession-level p-values were averaged by k-means group. **A**) Principal components 1 & 6 Core projections; individual average p-value across k-means group V; density plot of average p-values for group V. **B**) Principal components 1 & 7 Core projections; individual average *p*-value across k-means group II; density plot of average p-values for group II. **C**) Principal components 1 & 8 Core projections; individual average *p*-value across k-means group II; density plot of average *p*-values for group II. **D**) Principal components 1 & 9 Core projections; individual average *p*-value across k-means group III; density plot of average *p*-values for group III. **E**) Principal components 1 & 10 Core projections; individual average *p*-value across k-means group III; density plot of average *p*-values for group III.
