## Supplementary Figure S6 for "Multiple streams of genetic diversity in Japonica rice"

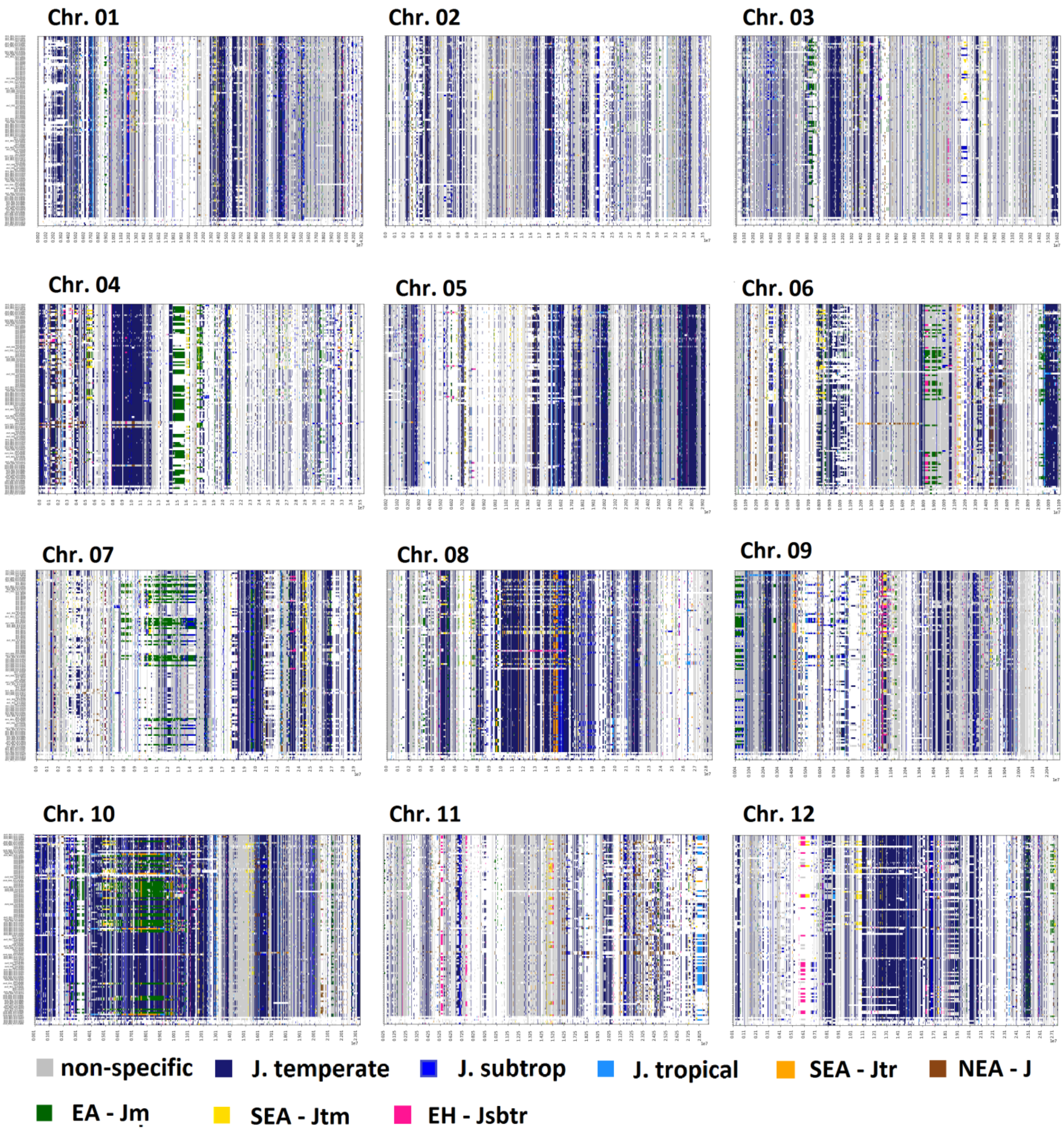


**Fig. S6 MAC assignment to groups indicative of Japonica substructure along the genome of tropical Japonica accessions.** Vectors of local cluster p-values were extracted for significance of pure Japonica assignments across tropical Japonica genomes. Vectors were grouped following PCA using k-means clustering at K = 10. Average group profiles were studied and related to Japonica sub-taxa. Local haplotypes were analysed for maximum affinity to clustered vectors and classified accordingly.
