## Supplementary Figure S7 for "Multiple streams of genetic diversity in Japonica rice"

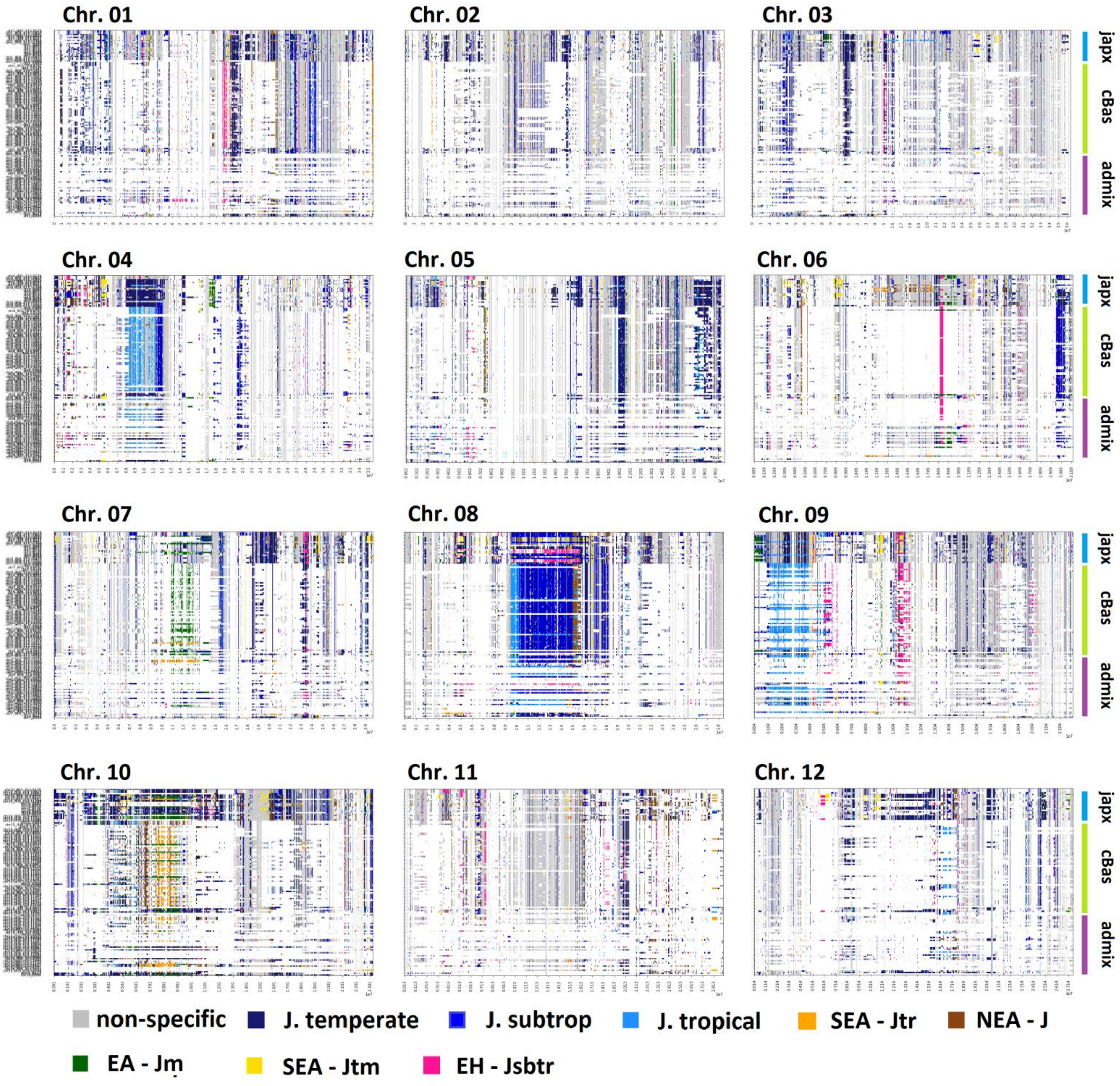


**Fig. S7 MAC assignment to groups indicative of Japonica substructure along the genome of tropical admixed accessions.** Vectors of local cluster p-values were extracted for significance of pure Japonica assignments across tropical japx, *cBasmati* and admix genomes (Wang *et al.* 2018). Vectors were grouped following PCA using k-means clustering at K = 10. Average group profiles were studied and related to Japonica sub-taxa. Local haplotypes were analysed for maximum affinity to clustered vectors and classified accordingly.
